# Modular platform for therapeutic drug delivery using trifunctional bio-orthogonal macromolecular conjugates

**DOI:** 10.1101/2025.07.24.666613

**Authors:** Danmeng Luo, Ning Wang, Hannah Major-Monfried, John Ralls, Sophia Rha, Stacy A. Maitland, Karthikeyan Ponnienselvan, Makiko Yamada, Daniel E. Bauer, Scot A. Wolfe, Alex Kentsis

## Abstract

Targeted delivery of macromolecular therapeutics holds great promise for overcoming the limitations of conventional small molecules, enabling modulation of protein-protein interactions and precise genome editing. However, efficient, safe, and cell type-specific delivery remains a major challenge. To address this, we developed a modular platform for synthesizing heterotrifunctional bio-orthogonal macromolecular conjugates (BMCs) by engineering diverse combinations of targeting ligands, cell-penetrating peptides (CPPs), and bioactive cargos. We optimized facile bioconjugation chemistries to generate BMCs with improved yields, structural integrity, and activity. Modular BMCs accommodate diverse components, including antibodies and receptor ligands for targeting, CPPs for intracellular trafficking, and optical probes, therapeutic peptidomimetics, and CRISPR-Cas9 nuclease as cargos to confer specific biological activities. We assayed their utility across multiple applications: BMCs with fluorescently labeled cargo revealed endosomal escape and intracellular accumulation; peptidomimetic MYB transcription factor inhibitor BMCs exhibited potent anti-leukemic activity against acute myeloid leukemia cells; and Cas9 BMCs achieved rapid delivery and cell type-specific gene editing in human cells. The BMC approach enables customizable delivery of functional macromolecules, nominating BMCs as a broadly applicable platform for biomedical applications.

**One-Sentence Summary:** The establishment of modular platform for synthesizing bio-orthogonal macromolecular conjugates (BMC) enabled fast and targeted delivery of membrane-impermeant macromolecular drugs.

## Introduction

The therapeutic potential of proteins and polypeptides—ranging from enzymes and antibodies to growth factors and transcription modulators—has sparked widespread interest in their clinical applications (*1–4*). Protein therapeutics exhibit distinctly high specificity and affinity to their biological targets, which are frequently “undruggable” by traditional synthetic small molecule drugs (*5–8*). Furthermore, protein therapeutics are typically biodegradable and less likely to generate toxic metabolites, contributing to a more favorable safety profile. Their large surface areas and structural molecular diversity also allow for the design of tailored therapeutics that can be engineered with desired pharmacokinetic properties and conditional activation in tissues (*9*, *10*).

However, efficient intracellular delivery of these generally membrane-impermeable macromolecules remains a formidable challenge. A variety of protein delivery systems have been developed, including liposomes, polymeric nanoparticles, viral vectors, and protein-based carriers, to enhance cellular uptake or improve pharmacokinetic bioavailability (*11*, *12*). Despite significant progress, current protein delivery platforms face several critical bottlenecks. In particular, poor selectivity and non-specific uptake by off-target tissues can lead to systemic toxicity and reduced bioavailability at the intended therapeutic sites of action, and thus often compromise the therapeutic efficacy of protein therapeutics.

To overcome these limitations, several bioengineering strategies have been explored to improve the selectivity of protein therapeutics. Incorporating targeting ligands into protein therapeutics offers a strategic advantage by enhancing tissue and cell type specificity. These ligands bind selectively to surface receptors expressed on diseased cells and facilitate receptor-mediated endocytosis, resulting in reduced off-target effects and enhanced cellular uptake (*13*). While targeting ligands have been conventionally assembled with cytotoxic drugs in the form of antibody-drug conjugates to convey targeted delivery of small molecule toxins (*14–16*), their direct use for protein therapeutics has not been systematically explored. A key barrier in this approach is the absent or hindered endosomal escape of macromolecular drugs upon internalization through receptor-based endocytosis. As endosomal escape is required for biological activity of therapeutics targeting nuclear or cytoplasmic compartments (*17*), approaches to facilitate endosomal escape with minimal cellular toxicity are required.

Cell-penetrating peptides (CPPs) are relatively short peptides with distinct physicochemical properties that have emerged as effective tools to enhance the intracellular delivery of protein therapeutics (*18*). CPPs such as TAT, penetratin, and R9 have been widely studied for their ability to transport various macromolecules—including enzymes, antibodies, and transcription factors—into the cytosol or nucleus of target cells (*19–23*). Recent advances have focused on designing novel CPPs with enhanced membrane penetrating capabilities and reduced cytotoxicity (*24*, *25*). Although their mechanisms of cell entry are not fully understood, transient membrane disruption is thought to be required for the penetration and subcellular translocation. Indeed, we and others have recently reported improved CPPs with enhanced nuclear and cytoplasmic delivery exceeding hundreds of millions of molecules per human cell while maintaining minimal membrane disruption and negligible toxicity *in vitro* (*24*).

Here, we report a modular synthetic platform for assembling macromolecular protein therapeutic cargos with both targeting ligands and CPPs in the same construct to achieve carrier-cargo-CPP trifunctional bio-orthogonal macromolecular conjugates (BMCs). Using bio-orthogonal synthetic chemistry with widely available reagents, we investigated three different molecular engineering strategies to assemble diverse BMCs with high yield, purity, and biological activity. We show that the modular BMCs platform can incorporate diverse targeting ligands, CPPs, and functional cargos—including optical probe for subcellular imaging, therapeutic peptidomimetic for modulating transcription factors, and CRISPR-Cas9 nuclease for therapeutic gene editing—to render specific biological activities. These results nominate BMCs as a broadly accessible and versatile approach for cell type-specific intracellular delivery of bioactive macromolecules for biomedical applications.

## Results

### Modular engineering of bio-orthogonal macromolecular conjugates (BMCs)

First, we sought to design a modular strategy for engineering and synthesizing macromolecular conjugates that would enable cell type-specific delivery, efficient intracellular internalization and sufficient endosomal escape of bioactive macromolecules. We envisioned the tripartite BMCs, containing i) targeting ligand such as antibody or receptor ligand (eg. plasma protein transferrin) to serve as carrier for targeted delivery, ii) CPP to confer membrane translocation and endosomal escape, and iii) macromolecular cargo to execute designed intracellular activity (**Fig. 1A**). To enable specific and efficient synthesis, we used bio-orthogonal coupling chemistry, including amines reacting with *N*-hydroxysuccinimide (NHS) esters, thiols reacting with maleimides, asymmetric disulfide bond formation, and strain-promoted azide-alkyne cycloaddition (SPAAC). These reactions were selected due to their high efficiency under mild native conditions, which enabled the preservation of the structure and function of bioactive macromolecules (**Fig. 1B**).

**Figure 1.**
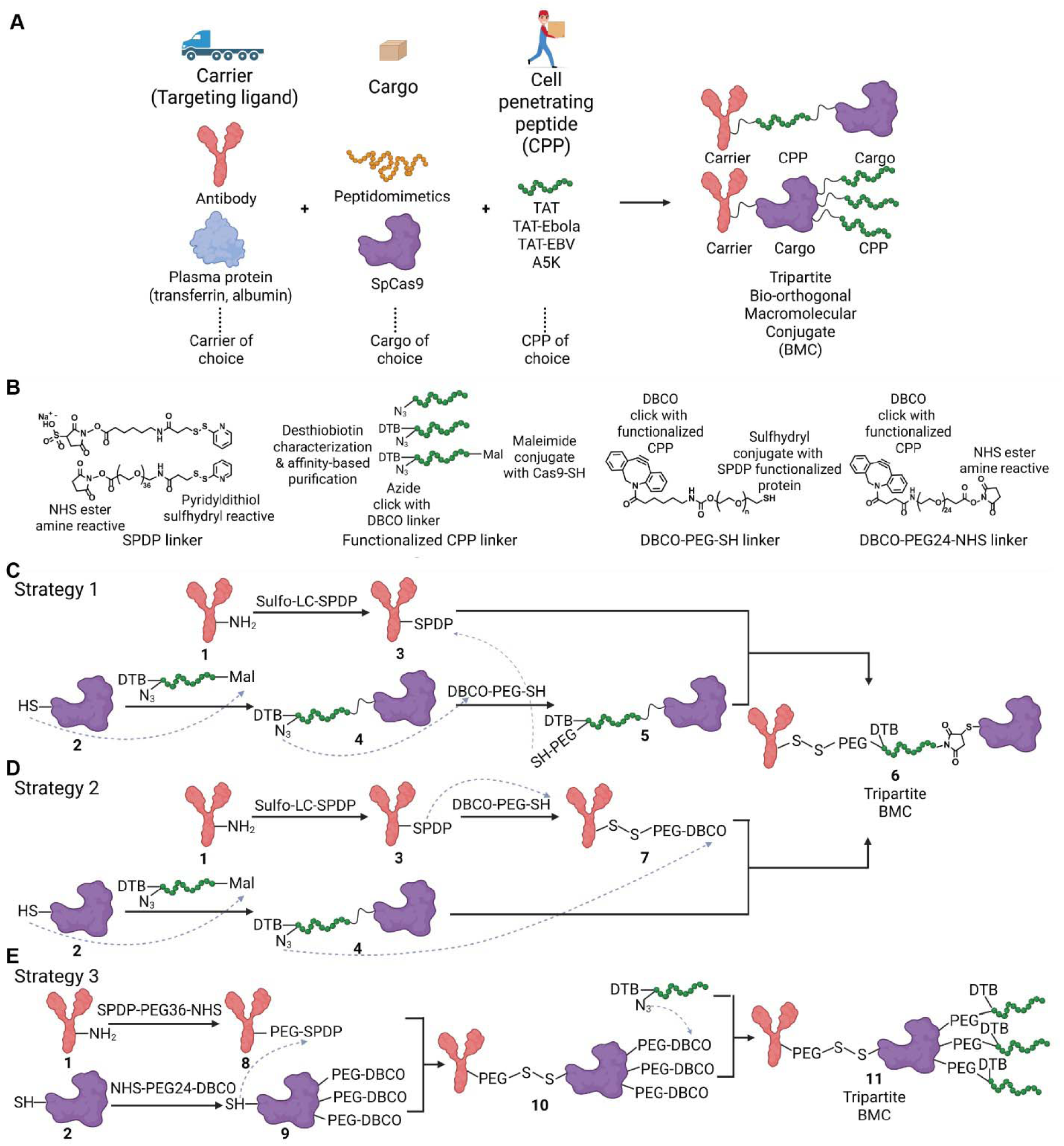
Schema for the design and synthesis of bio-orthogonal macromolecular conjugates (BMCs). (**A**) Graphic illustration of the BMC platform for therapeutic macromolecule delivery. (**B**) Key reagents used in assembling tripartite BMCs. All essential functional groups for bio-orthogonal conjugation are labeled with their use in the corresponding reactions. (**C**lll**E**) Synthetic strategies 1, 2, and 3 to construct tripartite BMCs. Arrows with dashed line represent reactions occurring between the pointed functional groups.

To assemble diverse BMC architectures with different structural features, we investigated three synthetic strategies to assemble tripartite BMCs (**Fig. 1C**lll**E**). These three strategies differed by module assembly sequence and were distinguished by the last module assembled into the BMCs, enabling efficient construction and screening of BMCs optimized for specific biological activities. Namely, **Strategies 1**, **2**, and **3** can be used to define the functional contribution of specific carriers, cargos, and CPPs, respectively. Importantly, we designed these strategies in a convergent manner and established a concise three-step modular conjugation scheme to produce the final BMC products (**Fig. 1C**lll**E**). To enable this, we designed CPPs with three different functional groups, i) maleimide to react with the cysteine side-chain embedded in the cargo protein, ii) azide to enable the bio-orthogonal SPAAC click chemistry, and iii) desthiobiotin (DTB) functional group to enable quantitative validation of conjugation efficiency and/or affinity-based purification, based on desthiobiotin’s specific binding to streptavidin and elution with biotin (*26*), respectively (**Fig. 1B**).

For **Strategies 1** and **2** (**Fig. 1C** and **D**), BMC synthesis proceeds with thiolation of the carrier (**1**) using sulfosuccinimidyl 6-(3’-(2-pyridyldithio)propionamido)hexanoate (sulfo-LC-SPDP) linker to form the carrier-SPDP intermediate (**3**). In parallel, specific cargo (**2**) is reacted with the tri-functionalized CPP via thiol-maleimide reaction to produce the cargo-CPP intermediate (**4**). **Strategies 1** and **2** differ in the specific orientation of the dibenzocyclooctyne-PEG-thiol (DBCO-PEG-SH) linker, which can be conjugated to the cargo versus the carrier, respectively (**Fig. 1C** and **D**). For **Strategy 1**, the DBCO-PEG-SH linker is reacted with the cargo-CPP intermediate (**4**) using SPAAC click chemistry to generate the cargo-CPP-SH (**5**), which is then conjugated with the carrier-SPDP intermediate (**3**) via asymmetric disulfide bond formation to generate the final tripartite BMC (**6**). For **Strategy 2**, the DBCO-PEG-SH linker is attached to the carrier-SPDP intermediate (**3**) via asymmetric disulfide bond formation, and the resultant carrier-DBCO (**7**) is then assembled with the cargo-CPP (**4**) via SPAAC click chemistry to form the final tripartite BMC (**6**). Both **Strategies 1** and **2** enable site-specific installation of the carrier-CPP-cargo tripartite BMC, with explicit control of cargo valency for functional and therapeutic applications.

In contrast, **Strategy 3** leverages the initial modification of the cargo (**2**) with an NHS-PEG24-DBCO linker to install a DBCO functional group as a handle for bio-orthogonal CPP conjugation at the final step (**Fig. 1E**). The resultant cargo-DBCO intermediate (**9**) is directly conjugated to a thiolated carrier (**8**) through asymmetric disulfide bond formation. Lastly, the obtained carrier-cargo-DBCO intermediate (**10**) is then assembled with an azide-functionalized CPP to construct the final tripartite BMC (**11**). Importantly, **Strategy 3** provides enhanced versatility for screening different CPPs and enables variable CPP valency by controlling the stoichiometry of the NHS-DBCO handle installation.

To investigate the potential versatility and generalizability of the designed BMC synthesis strategies, we chose to test several different cargo, carrier, and CPP components. For cargos, we tested two candidate macromolecules with varying size and application. As CRISPR-Cas9 system is revolutionizing the field of gene editing, its cell type-specific delivery is highly desirable for biomedical research and therapeutic applications (*27–29*). In addition, as a 160-kDa protein, Cas9 represents large macromolecular cargo with stringent requirement for accurate folding and structural integrity to confer specific activities (*30*, *31*). As a challenging cargo, delivery of Cas9 has been achieved mostly as nanoparticle (nanocluster, nanocomplex, etc) format, while most of them lacked specificity to defined cell populations and often accumulated in liver when administered (*27–29*, *32*, *33*). Similarly, intracellular protein-protein interactions are a central mechanism of biological signaling, while many of these interactions involve targets frequently considered ‘undruggable’ with small molecule therapeutics (*6*, *34–36*). Here, we selected the 5-kD selective peptidomimetic inhibitor of the MYB:CBP/P300 transcription coactivation complex, termed CRYBMIM, which specifically blocks oncogenic gene expression in leukemia cells while relatively sparing normal hematopoietic progenitor cells (*37*, *38*). Importantly, both Cas9 and CRYBMIM have limited cell penetrating activities on their own, necessitating improved strategies for efficient intracellular delivery.

Furthermore, to pursue cell type-specific or broad tissue delivery, we evaluated various targeting ligands as carriers, including immunoglobulin antibody and blood plasma carrier proteins transferrin and albumin, given their established use in current medical practice. For antibody, we specifically selected one targeting the cell surface receptor KIT, given its specific expression by blood stem and progenitor cells, which are important targets for therapeutic gene editing approaches currently being explored to correct monogenic blood disorders, such as sickle cell disease (*39–41*).

Lastly, we utilized structurally diverse CPPs, including the prototypical cationic CPP TAT, as well as recently developed improved CPPs, TAT-Ebola, TAT-EBV and A5K (**Table S1**)(*24*, *42*). These CPPs represent different physicochemical classes, including cationic, amphipathic, and chimeric penetrator molecules. In all, this battery of cargos, carriers, and CPPs constitutes a varied set of components with distinct chemical and structural properties intended to elucidate the structure-activity principles of the BMC approach.

### BMCs enable enhanced biological activity of therapeutic macromolecules

First, we used fluorescently-labeled peptidic molecule as cargo and transferrin as carrier to establish the initial macromolecular conjugation methods. To monitor the intracellular internalization of macromolecular conjugates, we developed a dual fluorophore labeling system to specifically label cargo and carrier molecules, thereby enabling their analysis in live human acute myeloid leukemia (AML) cells using confocal fluorescence emission microscopy (**Fig. 2A** and **S1**). We first labeled transferrin with a near-infrared fluorophore DyLight800 using NHS-amine chemistry, followed by thiolation using sulfo-LC-SPDP. The resultant Tf-DyLight800-SPDP was then reduced using tris(2-carboxyethyl)phosphine (TCEP) to reveal the thiol, which was then reacted with TAT cargo peptide containing fluorescein amidite (FAM) and a cysteine residue protected with 3-nitro-2-pyridinesulfenyl (Npys), Cys(Npys)-TAT-FAM (**Table S1**), via asymmetric disulfide bond formation to assemble the final transferrin-DyLight800-TAT-FAM BMC.

**Figure 2.**
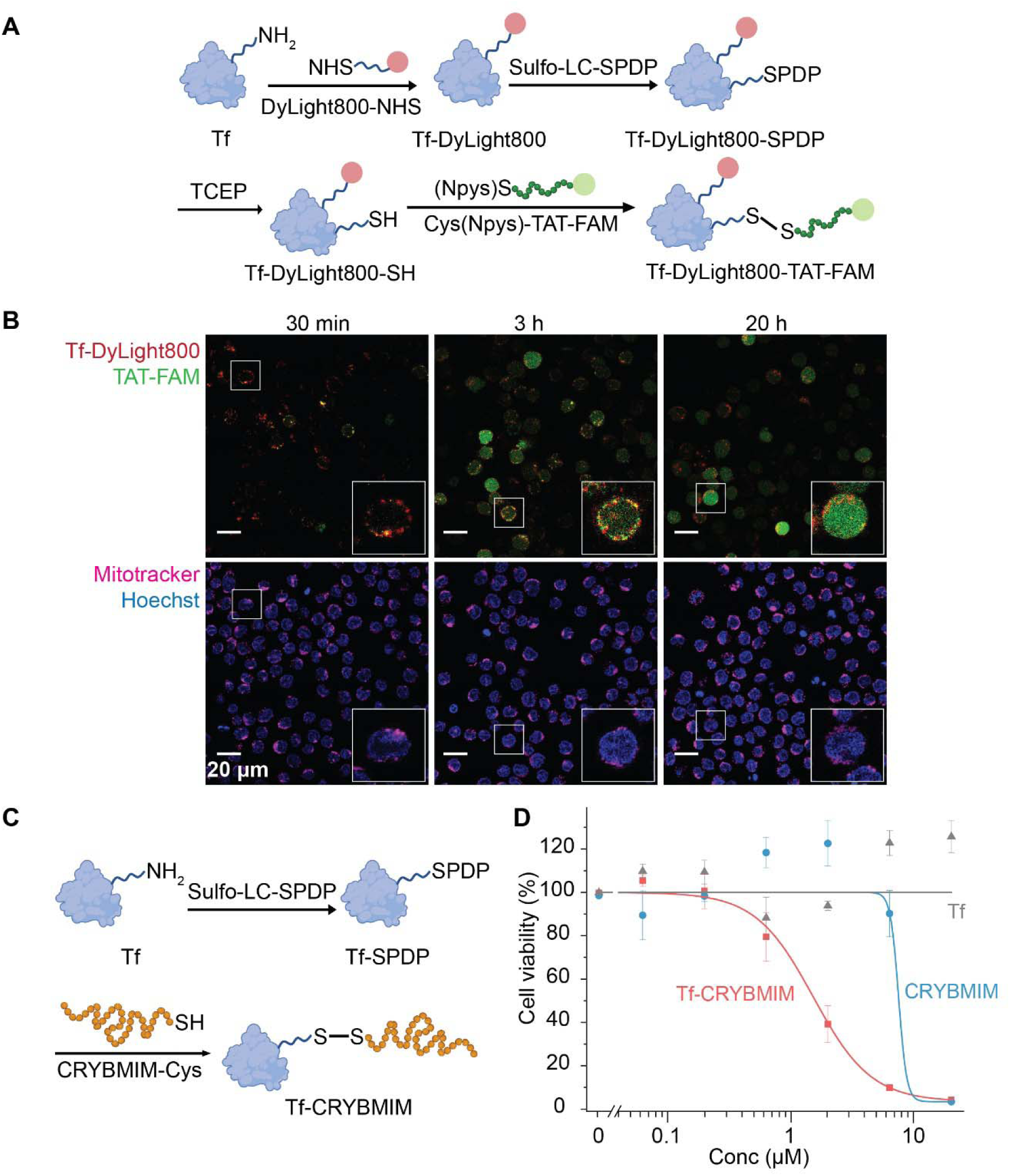
Efficient intracellular delivery of MYB mimetic inhibitor cargos with transferrin BMCs to target AML. (**A**) Synthetic strategy for constructing Tf-DyLight800-TAT-FAM BMC. Red and green spheres denote DyLight800 and FAM fluorophore, respectively. (**B**) Time-dependent intracellular internalization of Tf-DyLight800-TAT-FAM BMC and subsequent cargo release in MV-411 AML cells. Representative live cell confocal fluorescence emission microscopy photographs were shown at 30 minutes, 3 hours, and 20 hours after 40 nM Tf-DyLight800-TAT-FAM BMC treatment. Mitotracker (magenta) was used to define live cells and Hoechst dye (blue) was used to define nuclei, with TAT-FAM shown in green and DyLight800 shown in red. Bottom right insets show magnification of representative individual cells. Scale bar = 20 μm. (**C**) Synthetic strategy for constructing Tf-CRYBMIM BMC. (**D**) Anti-leukemic potency of Tf-CRYBMIM BMC (red squares), comparing with CRYBMIM (blue circles) and Tf alone (gray triangles) using MV-411 AML cells treated for 5 days, with cell viability monitored using CellTiter-Glo luminescence, normalized to cells treated with PBS control. Symbols and whiskers show means and standard deviations for three biological replicates. Tf stands for transferrin.

Multi-color fluorescence emission live cell confocal microscopy revealed time-dependent intracellular internalization of transferrin-DyLight800-TAT-FAM BMC in cultured MV-411 human AML cells, followed by subsequent cargo release and nuclear accumulation (**Fig. 2B**). Intracellular internalization of transferrin BMCs demonstrated that the modified transferrin carrier remained competent for binding and endocytosis via cell surface transferrin receptors. We also confirmed that the observed intracellular accumulation was not due to cellular toxicity or non-specific membrane rupture, as cells exhibiting BMC internalization also had normal mitochondrial activity, as measured using the MitoTracker Red FM dye (magenta), a specific fluorescent probe of cellular mitochondrial activity (*43*). Similarly, we confirmed nuclear accumulation of BMC-delivered Cys-TAT-FAM cargo (**Fig. 2B**), as identified by co-localization with nuclear chromatin, as measured with the membrane-permeable fluorescent DNA-intercalating Hoechst 33342 dye (blue).

Likewise, we also validated that the disulfide-conjugated BMC cargo was competent for endosomal reduction and release, likely conferred via the membrane destabilizing activity of TAT CPP (*20*, *21*, *23*). As indicated in **Fig. 2B**, puncta could be observed within 30 minutes of BMC treatment with overlapping fluorescence emission (yellow) from transferrin DyLight800 (red) and cargo CPP FAM (green), demonstrating cellular internalization of intact BMCs. Notably, after 3 hours of BMC exposure, Cys-TAT-FAM cargo began to appear separately from the transferrin carrier signals and diffusely throughout the cytosol and nucleus. Ultimately, after 20 hours of BMC exposure, no cargo and carrier colocalized puncta were observed, consistent with complete release and nuclear delivery. Thus, transferrin BMCs enabled endosomal escape and intracellular cargo delivery to live human cells.

Having confirmed the intracellular delivery of transferrin BMCs, we synthesized transferrin BMCs containing the peptidic MYB inhibitor CRYBMIM (**Fig. 2C** and **S2**). We have validated the on-target activity of CRYBMIM to suppress MYB-dependent oncogenic gene expression in AML cells using biochemical disassembly of the MYB:CBP chromatin complex, suppression of MYB-dependent gene expression using transcriptomics, and on-target anti-leukemia activity using functional genetic resistance studies (*37*, *38*). We synthesized CRYBMIM containing 3-nitro-2-pyridinesulfenyl (Npys)-protected terminal cysteine residue (**Table S1**) and revealed the thiol immediately before conjugation using reduction with TCEP. Then, we thiolated transferrin using the aforementioned SPDP linkage, and reacted CRYBMIM-Cys under oxidizing conditions via asymmetric disulfide bond formation to form the transferrin-CRYBMIM BMC. Treatment of MV-411 leukemia cells with transferrin-CRYBMIM BMC demonstrated more than four-fold improvement in anti-leukemia potency as compared to free CRYBMIM and measured using CellTiter-Glo luminescent cell viability assay (*IC_50_* = 1.6 ± 0.17 versus 7.1 ± 0.50 µM, respectively; t-test *p* = 4.4E-3; **Fig. 2D**). Thus, the BMCs strategy is able to enhance the biological activity of therapeutic macromolecules, due to improvements in their intracellular delivery.

### Convergent construction of tripartite BMCs with large macromolecular cargos

To investigate the utility of BMC engineering for cell type-specific delivery of large macromolecular cargos, we synthesized Cas9 tripartite BMCs using all three designed strategies (**Fig. 1C**lll**E**). We used Cas9 proteins engineered with three nuclear localization signals (termed 3xNLS-SpCas9) as the initial template, based on prior work that optimized this construct for intracellular gene editing activity by electroporation (**Fig. 3A**)(*44*). We reasoned that the activity of Cas9 BMCs may depend on the position of tethering of the carrier to the Cas9 nuclease, given its requirement for complex formation with a guide RNA (gRNA) to identify its target site and the necessity of conformational reorganization of its nuclease domains upon DNA target site licensing to achieve DNA cleavage (*30*, *31*, *45–47*). Thus, based on the atomic-resolution structure of the gRNA and DNA-bound Cas9 (*31*, *46*, *48*), we engineered three different 3xNLS-SpCas9 variants (V1, V2 & V3) containing single cysteine residue at different positions within Cas9, likely to be amenable to macromolecular tethering based on prior fluorescent probe coupling studies or avoid interference in binding with the gRNA and DNA substrate (**Fig. 3A** and **B, Tables S2** and **S3**) (*45*, *46*).

**Figure 3.**
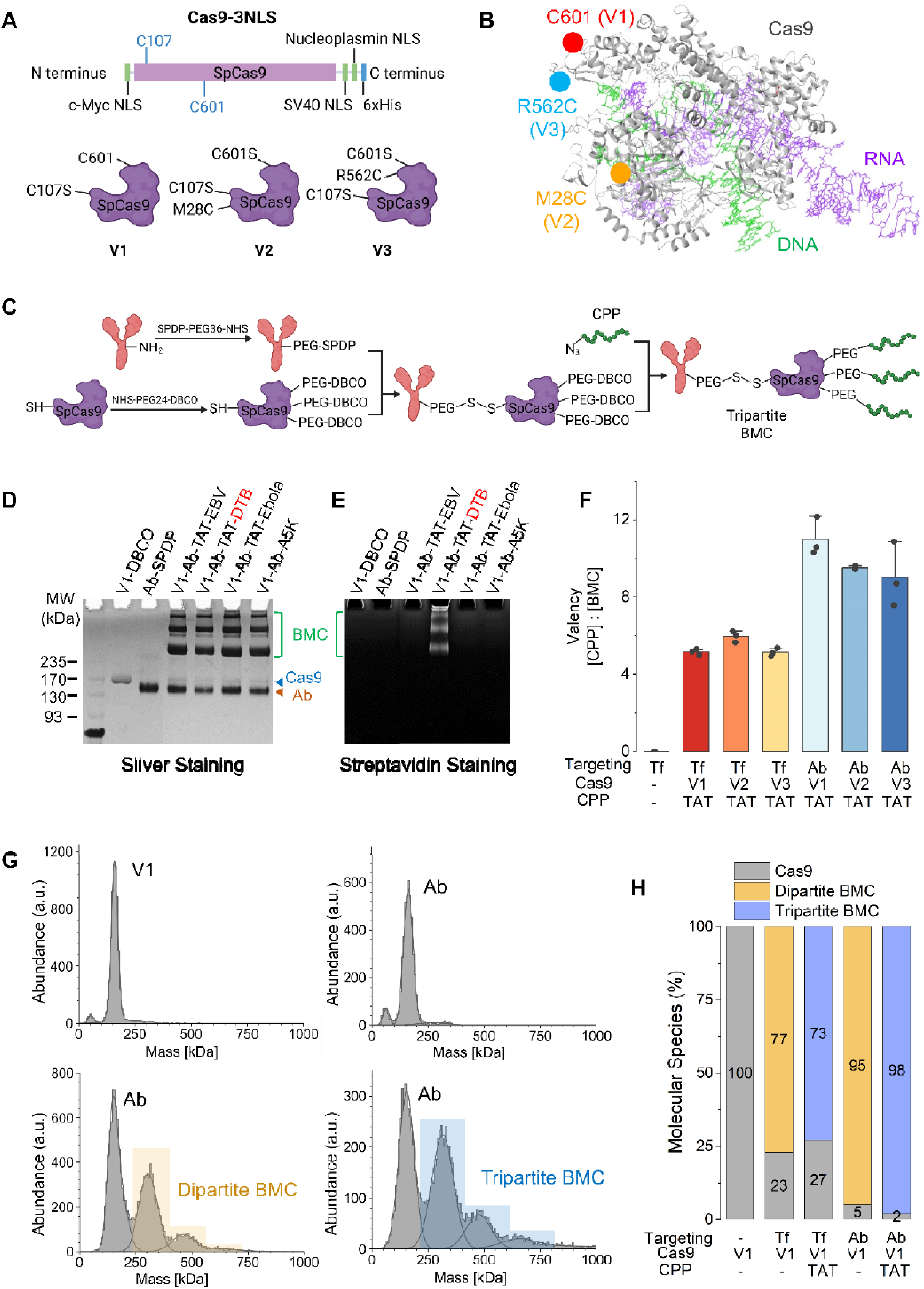
High yield synthesis of Cas9 tripartite BMCs. **(A)** Design schematic of the 3xNLS-SpCas9 nuclease, and its variants (V1, V2, and V3), showing three nuclear localization signals (NLS), 6xHis purification tag, and native C107 residue. (**B)** Structural model of Cas9 nuclease (gray), with its gRNA (purple) and DNA (green) substrates, based on the PDB 4oo8 structure, highlighting V1 (red), V2 (yellow), and V3 (blue) amino acid substitutions. (**C)** Synthesis strategy 3 used for engineering Cas9 tripartite BMCs. (**D and E)** Analysis of the reaction intermediates (Cas9 and Ab) and final products (BMC) using SDS-PAGE with silver staining (D) and streptavidin staining (E). TAT peptide functionalized with desthiobiotin is labeled in red. (**F)** Quantitation of desthiobiotin moieties in Cas9 BMCs to estimate CPP valency, based on the molar ratio of CPP and BMC for BMCs with V1, V2 or V3 Cas9 as cargo, Tf or Ab as carrier, and TAT as CPP. Serial dilution of stock solution of desthiobiotin TAT peptides was used as reference standard to establish calibration curves, with the transferrin reaction intermediate (Tf-SPDP) used as the negative control. Symbols and whiskers represent means and standard deviations of 3 replicates. (**G)** Mass photometry histograms of the reaction intermediates and final BMC products measured at 250 nM concentrations, showing relative abundance as a function of measured molecular mass, for the expected tripartite (mean mass=305 kDa), Cas9-Ab-Cas9 oligomer (mean mass=465 kDa), and tripartite BMC dimer (mean mass=610 kDa). (**H)** Quantitative analysis of the mass photometry data to estimate relative BMC yields, containing V1 Cas9 as cargo, Tf or Ab as carrier, and with or without TAT as CPP. Molecular binding events of tripartite, oligomer and tripartite dimer were combined in calculating the product yield based on the consumption of Cas9 instead of the antibody/transferrin, which were used in slight excess to Cas9 in the synthesis.

Accordingly, we used **Strategies 1** and **2** to construct Cas9 tripartite BMCs. First, tri-functionalized TAT CPP containing the N-terminal desthiobiotin and azide plus C-terminal maleimide functional groups, as well as the dual-functionalized DBCO-PEG-SH linker were incorporated to tether the cargo Cas9 with the carrier KIT antibody to form the tripartite BMC (**Fig. 1 C** and **D**). We confirmed the site-specific synthesis of the tripartite BMC using sodium dodecyl sulfate polyacrylamide gel electrophoresis (SDS-PAGE) followed by silver staining for protein analysis under denaturing conditions (**Fig. S3**), as well as quantitative mass photometry of BMC complexes in aqueous solution under native conditions (**Fig. S4**). We observed the reaction yields of tripartite BMC were modest with **Strategies 1** and **2** (mean 7 and 8%, respectively), which were not improved substantially by varying reaction stoichiometries, temperature, duration, buffer conditions, or linker rigidities and lengths (**Fig. S3**lll**S5**).

In contrast, with **Strategy 3** we achieved over 90% reaction yield in assembling Cas9 tripartite BMC, as evident from almost complete consumption of the starting Cas9 material and specific generation of high-molecular mass conjugates (**Fig. 3D** and **fig. S6**). We confirmed the correct assembly of the Cas9 tripartite BMCs using SDS-PAGE (**Fig. 3D**) and mass photometry (**Fig. 3G** and **H**), which demonstrated the generation of monovalent carrier:cargo BMC complexes with the mean molecular mass of 305 kDa, in comparison to the mean 160-kDa and 145-kDa mass of the reactants, corresponding to Cas9 cargo and KIT antibody carriers, respectively. We also observed higher order assemblies that likely represent the coupling of more than one Cas9 protein to the antibody.

To orthogonally validate the correct BMC assembly and the intended CPP installation, we leveraged the desthiobiotin-containing CPP design (**Table S1**), as it is particularly challenging to measure CPP conjugation solely by changes in electrophoretic mobility or mass photometric light scattering, given TAT’s 2 kDa molecular mass, corresponding to just 7% of 305-kDa total mass of the conjugate. Thus, we confirmed BMC incorporation of desthiobiotin-containing CPP, consistent with the intendent BMC design, as assessed by Western blotting using fluorescently-conjugated streptavidin (**Fig. 3E**).

With this established method, we synthesized V1, V2, and V3 Cas9-Ab-TAT tripartite BMCs, achieving reaction yields of 98, 97, and 95%, respectively, as estimated by SDS-PAGE silver stain (**Fig. S7**). Thus, BMC assembly occurs efficiently for cysteines located at all three different surface positions on Cas9, suggesting similar steric accessibility and chemical reactivity for tripartite BMC formation. Leveraging the desthiobiotin-containing CPP design (**Table S1**), we estimated the BMC valency of installed CPPs using colorimetric desthiobiotin quantitation assay with free TAT-DTB peptides as calibration reference standards and identified mean CPP valencies of 11, 10, and 9 for the V1, V2, and V3 variants of Cas9-Ab-TAT BMCs, respectively (**Fig. 3F**). Multivalent CPP conjugation may enable enhanced BMC membrane permeabilization and endosomal escape of macromolecule cargos.

To test the robustness and potential generalizability of the BMC strategy, we used two additional carriers, transferrin and albumin, to generate the respective Cas9 tripartite BMCs, and all constructs were achieved with high reaction yields (**Fig. 3H** and **S8**lll**S9**). This produced Cas9-Tf-TAT BMCs with TAT valencies of 5, 6, and 5, for the V1, V2, and V3 variants, respectively, consistent with the differences in accessibility of the DBCO handle (**Fig. 3F**). Likewise, we constructed BMCs containing different CPPs (TAT-Ebola, TAT-EBV and A5K) spanning different physicochemical classes (**Fig. 3D and S7, Table S1**). This included relatively hydrophobic CPPs with limited aqueous solubility such as TAT-Ebola and A5K, for which we implemented a buffer system containing Pluronic F-127 poloxamer based on our prior studies for native protein extraction (*30*, *31*, *49*) **(Fig. S10**). Overall, this effort enabled successful synthesis of diverse BMCs, which incorporated different cargos, carriers and CPPs, demonstrating the versatility and generalizability of this modular platform technology.

### BMCs maintain Cas9 nuclease activity for sequence-specific gene editing

To evaluate whether Cas9 retains native folding and function after BMC synthesis, we assessed the nuclease activity of its tripartite BMCs using synthetic DNA substrates *in vitro*, and human genome editing in cells (**Fig. 4A**). First, we used a linearized plasmid DNA containing a single Cas9 target site as substrate for *in vitro* DNA cleavage assay to determine the effects of BMC synthesis on Cas9 ribonucleoprotein (RNP) formation with its complementary single guide RNA (sgRNA) and the overall nuclease activity. Next, we electroporated Cas9 BMC RNPs into human Kasumi-1 Traffic Light Reporter (TLR) cells (*50*, *51*)—where sequence-specific genome editing induces expression of mCherry fluorescent marker—to quantify intracellular editing efficiency by fluorescence-activated cell scanning (FACS) and confocal fluorescence emission microscopy.

**Figure 4.**
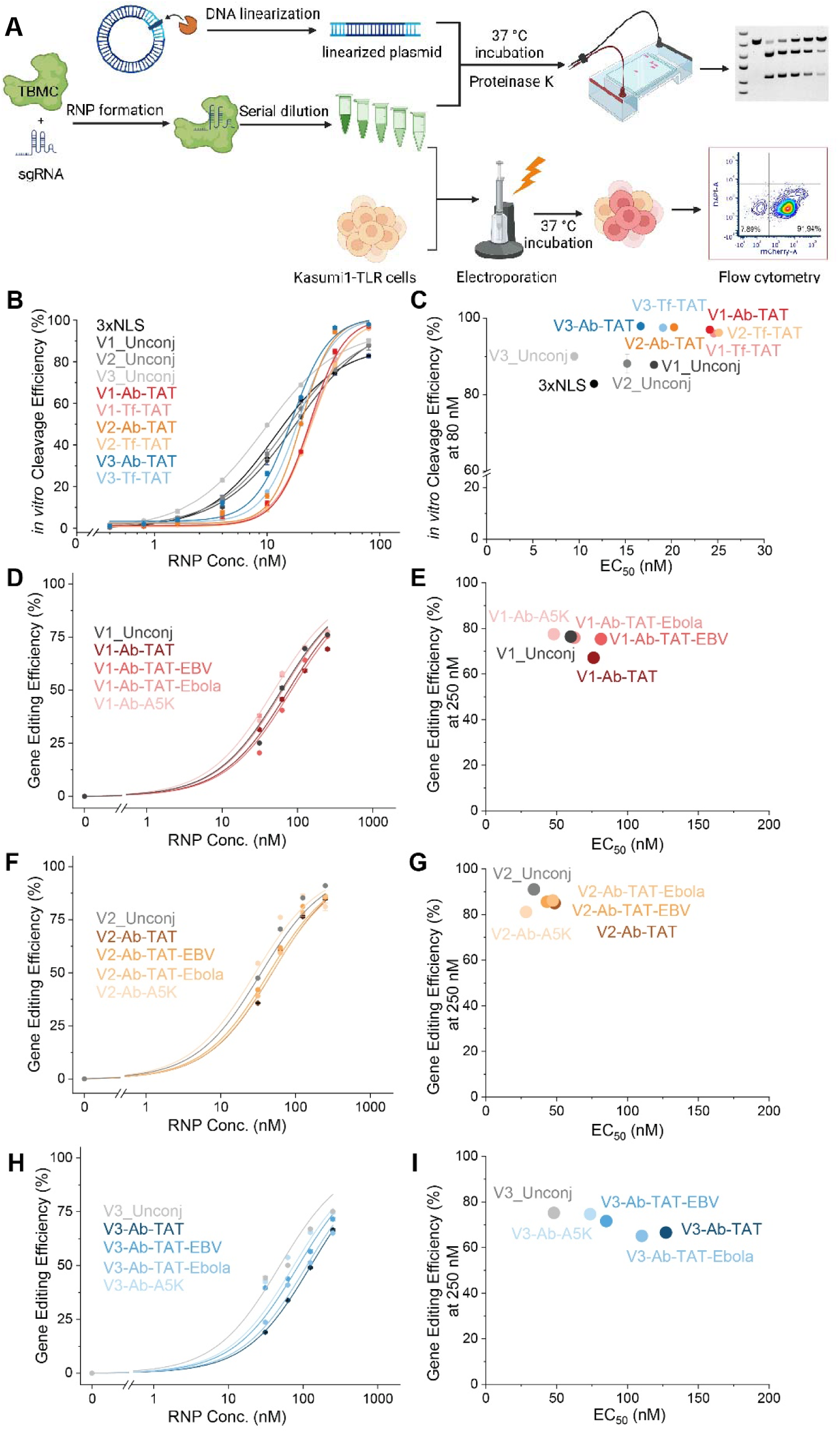
BMC bioengineering strategy preserves the bioactivity of Cas9 BMC cargos. **(A)** Schematic of the plasmid cleavage assay of BMC nuclease activity *in vitro* and upon electroporation in cells. (**B and C)** *In vitro* cleavage efficiency of Cas9 BMCs containing V1, V2, or V3 Cas9 variants as cargo, Tf or Ab as carrier, and TAT as CPP. The cleavage efficiency was assessed using gel electrophoresis analysis of linearized DNA cleavage as quantified by image densitometry, normalized to a DNA loading control. Unconjugated native Cas9 variants and 3xNLS-SpCas9 parent molecule were used as baseline control for comparison. (**D**lll**I)** Cellular BMC activity assessed upon electroporation in Kasumi-1 TLR cells and measured by FACS of mCherry-expressing cells. Cas9 BMCs containing V1 (D and E), V2 (F and G), or V3 (H and I) Cas9 variants as cargo, Ab as carrier, and TAT, TAT-EBV, TAT-Ebola, and A5K as CPP were tested. Unconjugated native Cas9 variants were used as baseline control for comparison. 3xNLS: 3xNLS-SpCas9; V1/V2/V3_Unconj: unconjugated native Cas9 variants. Data show 3 replicates.

We first validated that the editing efficiency of all three engineered V1, V2, and V3 Cas9 variants matched the parent 3xNLS-SpCas9 in apparent nuclease activity *in vitro* (**Fig. S11A**lll**E**). Additionally, we also demonstrated that the engineered Cas9 variant possessed adequate stability in serum-containing medium, cell lysates (**Fig. S11F** and **G**), and storage at 4°C (**Fig. S13**), as needed for BMC production for biomedical applications. We then tested V1, V2, and V3 Cas9-tripartite BMCs containing four different CPPs, TAT, TAT-EBV, TAT-Ebola and A5K, and two carriers, KIT antibody and transferrin. All BMC constructs retained enzymatic nuclease activity *in vitro* (**Fig. 4B** and **C, S12; Tables S4** and **S5**) and robust human gene-editing efficiency upon electroporation in cells (**Fig. 4D**lll**I**; **Table S6**). Interestingly, we observed differences between nuclease activities *in vitro* and in cells for distinct BMCs, suggesting that specific BMC configurations may have compensatory effects on RNP assembly *in vitro* with chromatin accessibility and cofactor activity in cells (**Fig. 4H** and **I; Fig. S12; Table S5-S6**). Similarly, conjugating a single highly cationic CPP TAT directly to Cas9 Cys via maleimide-thiol chemistry modestly reduced gene editing activity *in vitro* and upon electroporation in cells (**Fig. S14**). In contrast, their corresponding tripartite BMCs mostly maintained full biological activity upon electroporation in cells (**Fig. 4D**lll**G**; **Table S6**). These observations suggest that CPP-induced perturbations of Cas9 RNP stability can be apparently compensated by carrier conjugation within tripartite BMCs. Collectively, these findings demonstrate that the BMC synthetic approach preserves Cas9 conformational integrity and enzymatic nuclease activity in diverse tripartite BMCs.

### BMCs enable rapid and cell type-specific intracellular delivery of functional macromolecular cargos

To assess real-time intracellular delivery of Cas9 tripartite BMCs, we utilized DyLight488-labeled Cas9 constructs to treat Kasumi-1 TLR cells. Because Cas9 single-guide RNAs (sgRNAs) engineered for sequence-specific gene editing are 100 nucleotides in length, with only a small portion bound by Cas9 and the remainder solvent exposed, thus potentially interfering with the membrane penetration of Cas9 BMCs, we pre-loaded cells with sgRNA by electroporation, followed by washing to remove excess sgRNA, and treated sgRNA-loaded cells with Cas9 BMCs. We used gRNAs labeled with ATTO647 fluorophore to track their subcellular localization using time-lapse live cell confocal fluorescence emission microscopy. After just 30 minutes of exposure to Cas9 BMCs, where the excess BMCs were removed by cell washing, Cas9 BMC-treated cells displayed significantly more intracellular DyLight488 Cas9 puncta as compared to cells treated with Cas9 alone (mean 10 and 0.80 puncta/nucleus, respectively; one-way ANOVA with Tukey’s test *p* = 7.6E-4). In particular, BMC-treated cells exhibited specific co-localization between DyLight488 Cas9 puncta with both Hoechst-stained nuclear chromatin and ATTO647-labeled gRNA, confirming intended RNP assembly and nuclear import of BMC-delivered Cas9. Using time-lapse imaging, we observed time-dependent increase of nuclear accumulation of Cas9, culminating in successful gene editing to induce fluorescent mCherry expression upon 72 hours of culture, which is exclusively in BMC-treated cells (**Fig. 5D** and **E; Movie S1**). In contrast, unmodified Cas9 failed to generate any gene-editing reporter signal despite the observation of occasional intracellular puncta (**Fig. 5D; Movie S1**).

**Figure 5.**
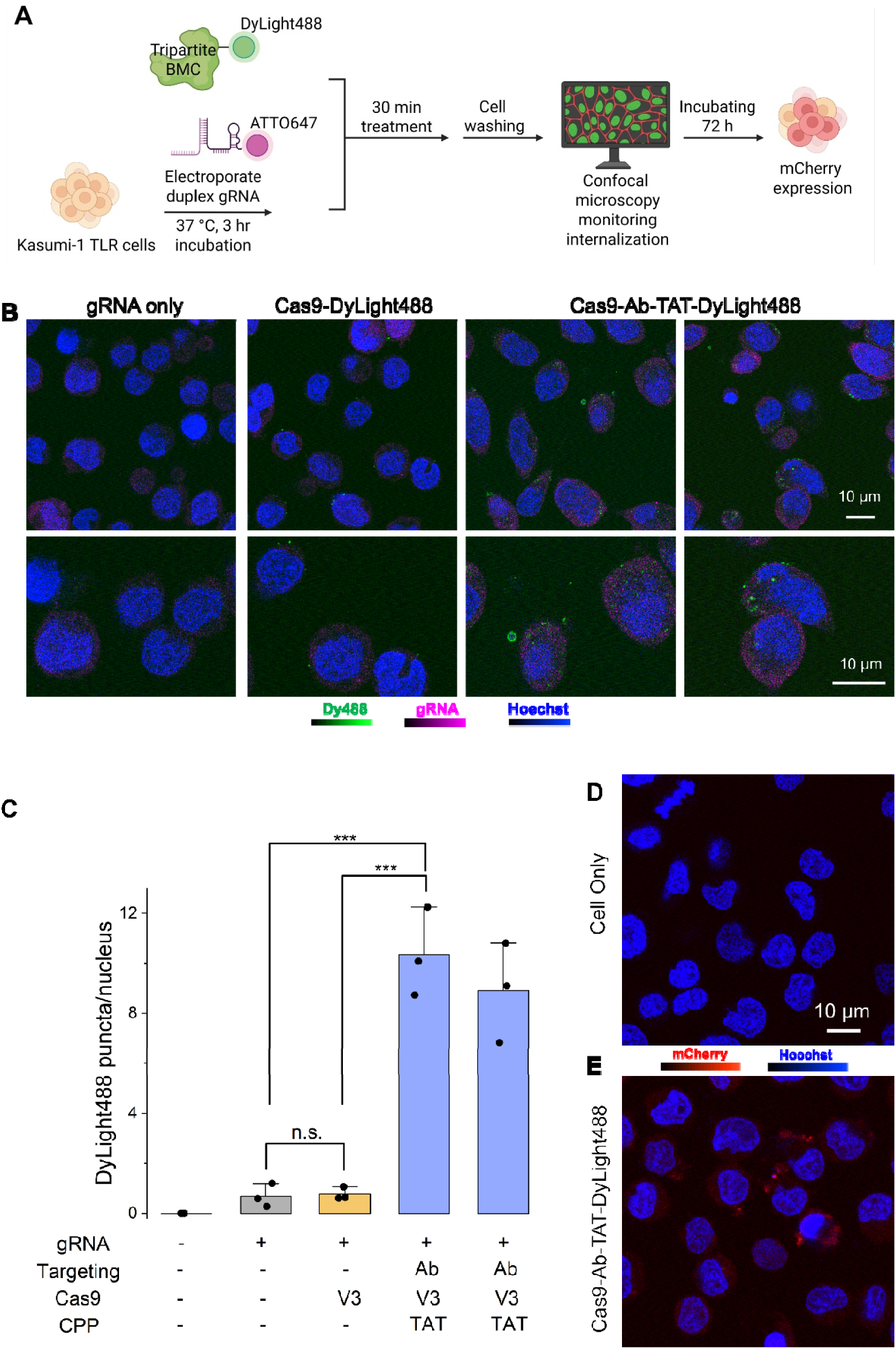
Rapid intracellular delivery of Cas9 tripartite BMCs. **(A)** Schematic workflow of experimental measurements of subcellular localization of BMC components using live cell confocal fluorescence emission microscopy. (**B)** Representative live cell confocal fluorescence emission microscopy photographs of the Kasumi-1 TLR cells treated with ATTO647-labeled gRNA (purple) only versus Cas9-DyLight488 (green) versus Cas9-Ab-TAT-DyLight488 BMC revealed the internalization of the fluorescently labeled Cas9 tripartite BMCs. Hoechst (blue) was used to define nuclei. z = 0.85 μm. Scale bar = 10 μm. (**C)** Quantitative image analysis of Cas9-DyLight488 puncta per Hoechst-stained nucleus of cells treated with PBS, gRNA only, Cas9-DyLight488, or Cas9-Ab-TAT-DyLight488 BMC. An average of 56, 46, and 25 nuclei were identified from the gRNA-only, Cas9-DyLight488 and Cas9-Ab-TAT-DyLight488 treated samples, respectively in triplicate measurements. Symbols and whiskers represent standard deviations of three technical replicates. (**D and E)** Representative live cell confocal microscopy photographs of cells 72-hours post Cas9 tripartite BMC treatment (E) in comparison of cell only (D). mCherry expression (red) was observed in Cas9 tripartite BMC treated cells, indicating successful nuclear delivery of functional Cas9 cargo, conferring sequence-specific gene editing to induce mCherry expression. Scale bar = 10 μm. * *p*≤0.05, ** *p*≤0.01, *** *p*≤0.001 for ANOVA Tukey’s test.

We next quantified sequence-specific BMC-induced gene editing in cells by measuring the ratio of mCherry-fluorescent cells via flow cytometry, which served as the ultimate readout of successful BMC nuclear delivery of functional Cas9 (**Fig. 6A**). We compared Cas9-Ab-TAT BMCs comprised of all three Cas9 variants and concentrations using 30 minutes of BMC treatment followed by cell washing to remove excess BMC. At 4 µM concentration, Cas9 V3 BMCs bearing TAT CPP achieved over 60% gene-editing efficiency as compared to 3.3% with unmodified Cas9 (two-way ANOVA with Tukey’s test *p* = 3.0E-7; **Fig. 6B** and **C**). Cas9 V3 variant consistently outperformed V1 and V2, suggesting that the respective cystine positions and conjugation sites can influence BMC internalization efficiency (**Fig. S15** and **S16**).

**Figure 6.**
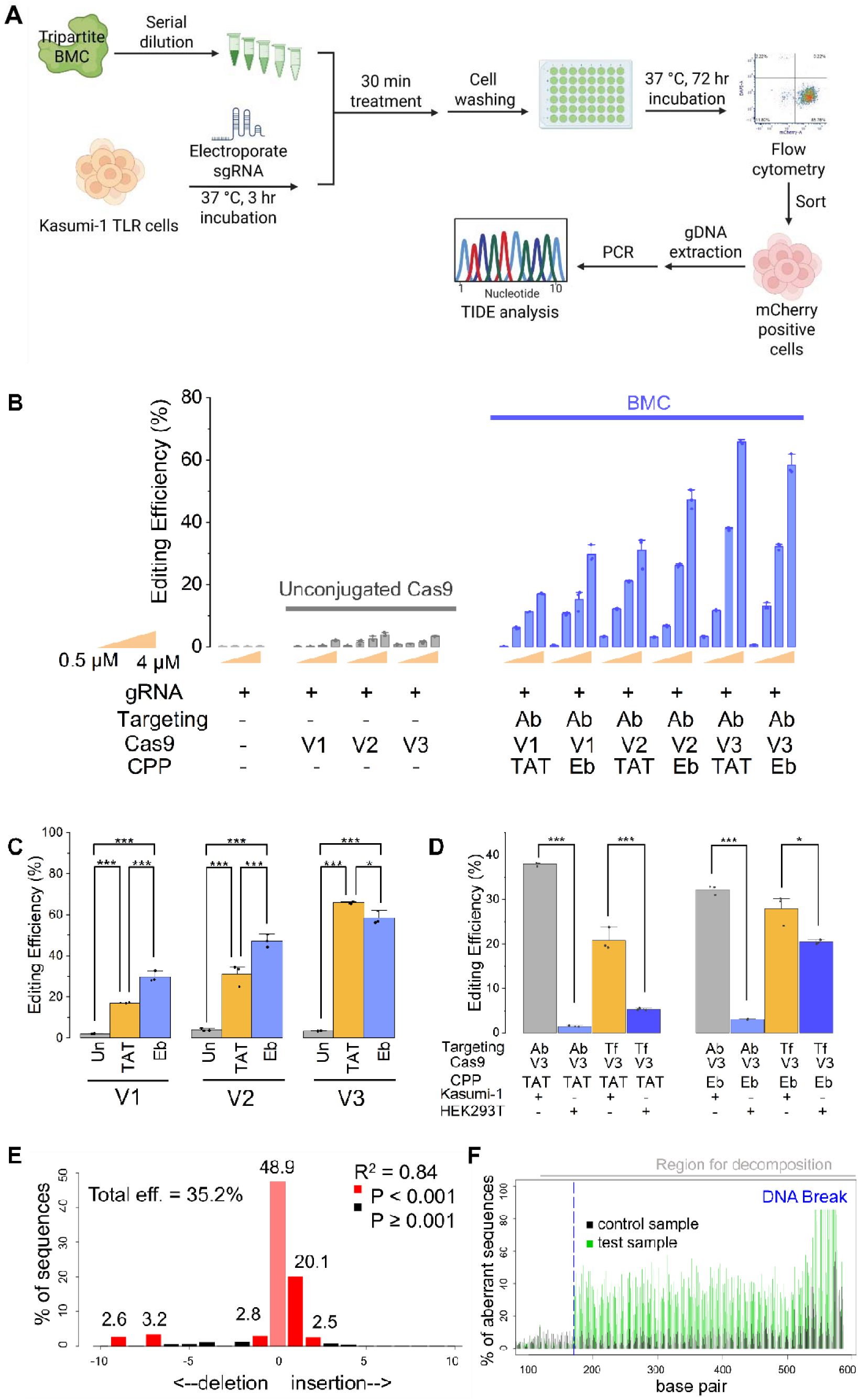
Cell type-specific human gene editing using Cas9 tripartite BMCs. **(A)** Schematic workflow of Kasumi-1 TLR or HEK393 TLR cells loaded with specific sgRNA by electroporation, followed by treatment with Cas9 tripartite BMCs and analysis of gene editing by flow cytometry and DNA sequencing. (**B)** Gene editing efficiency of unconjugated Cas9 (gray) versus BMCs (blue) in Kasumi-1 TLR cells as a function of increasing concentration for tested constructs. The Cas9 tripartite BMCs contained V1, V2, V3 Cas9 as cargo, KIT antibody (Ab) as carrier, and TAT or TAT-Ebola (Eb) as CPP. Bars and whiskers represent means and standard deviations of three biological replicates. (**C)** Editing efficiencies of 4 µM Cas9 tripartite BMCs in Kasumi-1 TLR cells. Cas9 tripartite BMCs contained V1, V2 or V3 Cas9 variants as cargo, KIT antibody (Ab) as carrier, and TAT (yellow) or TAT-Ebola (Eb, blue) as CPP. Unconjugated Cas9 variants (gray) were included as baseline control for comparison. Bars and whiskers represent means and standard deviations of three biological replicates. (**D)** Editing efficiencies assessed in Kasumi-1 TLR versus HEK293T TLR cells, treated with 2 µM Cas9 tripartite BMCs containing V3 Cas9 as cargo, transferrin (Tf) or KIT antibody (Ab) as carrier, TAT or TAT-Ebola (Eb) as CPPs. Bars and whiskers represent means and standard deviations of three biological replicates. (**E and F)** TIDE analysis of Sanger sequencing of TLR locus in BMC-treated mCherry-positive Kasumi-1 TLR cells, showing sequence-specific DNA editing. * *p*≤0.05, ** *p*≤0.01, *** *p*≤0.001 for ANOVA Tukey’s test.

Comparative analysis of TAT-Ebola, TAT-EBV, and A5K CPPs revealed that TAT-Ebola conferred improved human gene-editing activities across all Cas9 BMC variants (mean editing efficiency of V3 BMC at 4 µM of 58, 1.5, and 2.1%, respectively; one-way ANOVA with Tukey’s test *p* = 6.6E-6 and 6.7E-6 for TAT-Ebola versus TAT-EBV and A5K for V3 BMC, respectively; **Fig. 6B** and **C; Figs. S15** and **S16**).

To verify the accurate BMC gene-editing activity in cells, we purified mCherry-positive cells using fluorescence-activated cell sorting (FACS) and analyzed their genomic DNA (gDNA) using Tracking of Indels by Decomposition (TIDE) (*52*), which confirmed intended gene-editing at the targeted DNA position within the TLR locus with the total efficiency of 35.2% (*R^2^* = 0.84; **Fig. 6E** and **F**). Thus, both macromolecular cargo conjugation geometry and CPP identity represent important parameters for effective BMC-mediated intracellular delivery and Cas9 nuclease genome-editing activity in cells.

Finally, we investigated cell type-specific BMC delivery and carrier-dependent targeting effect by assessing BMCs with the specific targeting ligand KIT antibody versus the broad targeting ligand transferrin. We also utilized two different lineage human cell lines to test cell type specificity: Kasumi-1 cells with high KIT and transferrin receptor expression, whereas HEK293T cells with low KIT and moderate transferrin receptor expression. We analyzed four different Cas9 tripartite BMCs, with KIT antibody or transferrin serving as carriers, and TAT or TAT-Ebola serving as CPPs (**Fig. 6D** and **S17**). We found that the KIT antibody conferred strong specificity and enabled targeted delivery of the corresponding Cas9 tripartite BMCs, with V3-Ab-TAT and V3-Ab-TAT-Ebola tripartite BMCs specifically internalized in KIT-positive Kasumi-1 TLR cells as compared to KIT-negative HEK293T TLR cells (mean gene-editing efficiency of 38 and 33 versus 1.5 and 3.0%, respectively; one-way ANOVA with Tukey’s test *p* = 1.1E-7 and 1.1E-6 for V3-Ab-TAT and V3-Ab-TAT-Ebola, respectively; **Fig. 6D** and **S17**). In contrast, given the broader expression of its receptor, transferrin carrier BMCs were able to convey delivery into both cell types (21 and 28% versus 5 and 20% for V3-Tf-TAT and V3-Tf-TAT-Ebola BMCs in Kasumi-1 TLR and HEK293T TLR cells, respectively; **Fig. 6D** and **S17**). Thus, the carrier-dependent cell specificity, combined with modular flexibility in selecting CPP and cargos, demonstrates the versatility of the BMC platform and its potential for cell type-specific intracellular delivery of macromolecular therapeutics.

## Discussion

We have established a convergent, modular platform for synthesizing tripartite BMCs that integrates diverse macromolecular cargos—including small molecule optical probes, peptidomimetics, and macromolecular nucleases—with targeting ligands including antibodies and blood carrier proteins, and cell penetrating peptides conferring membrane permeabilization via a three-step bio-orthogonal synthetic strategy. In particular, the described **Strategy 3** consistently achieved greater than 90% synthetic yields across diverse cargo types, from short peptides to 160-kDa macromolecular enzymes. Biochemical and cellular assays confirmed that both inhibitory and catalytic macromolecular cargos retained biological activity upon BMC synthesis, while live-cell confocal microscopy imaging demonstrated rapid cytosolic and nuclear BMC macromolecular delivery on the timescale of minutes, yielding therapeutic functional effects including anti-cancer effects of peptidomimetic transcription factor inhibitor and sequence-specific genome editing by engineered Cas9 nuclease in human cells. Lastly, carrier choice of cell surface receptor-specific KIT antibody and blood plasma transferrin conferred cell type selective macromolecular delivery to receptor-expressing but not non-expressing cells.

While BMCs demonstrate robust performance *in vitro* and in cultured cells, their pharmacokinetics, biodistribution, and immunogenicity *in vivo* remain to be explored. Likewise, the current study relies on chemical linkers that may exhibit variable stability under physiological redox and proteolytic conditions. Although we observed efficient endosomal escape of BMC cargos, the exact mechanisms and potential side effect of membrane disruption by BMC-installed CPPs will require further study. While we derived the initial structure-activity relationships of tripartite BMCs with respect to the effects of linker length, CPP valency, and carrier density *in vitro*, their effects on tissue penetration and potential toxicities *in vivo* will need to be defined. Lastly, though the reported BMC synthetic strategy achieved over 90% yield, future studies will need to optimize this approach for large-scale manufacturing and scalability of multi-component BMCs for therapeutic and clinical-grade applications.

Nevertheless, the developed bio-orthogonal macromolecular conjugate (BMC) platform offers distinct molecular engineering features with important advances for cell type-specific delivery of membrane impermeable macromolecules, including those targeting protein-protein interactions or conferring enzymatic activities in various subcellular compartments in cells. In addition, adapting cleavable or stimuli-responsive linkers may enhance and control macromolecular cargo release while minimizing systemic exposures. Expanding the BMC cargo repertoire to include enzymes that reprogram metabolic pathways, peptidomimetics and proteolysis and proximity targeting chimeric molecules (PROTACs) that block and modulate intracellular signaling, synthetic scaffold proteins and transcriptional effectors among others, should broaden research applications and translational potential of BMCs. Likewise, coupling BMCs with imaging agents could enable theranostic applications, while combining multiple cargos may enable dynamic and synergistic therapies. Ultimately, the modular BMC platform offers a generalizable strategy for targeted intracellular delivery of diverse macromolecular drugs, with promising applications in gene therapy, oncology, metabolic disorders, and many other biomedical applications dependent on macromolecular modulators and therapeutics.

## Materials and Methods

### Reagents

The purified anti-human CD117 (KIT) antibody was obtained from Biolegend (cat. 313202) in azide-free phosphate-buffered saline (PBS). The human *apo*-transferrin (cat. T1147) and bovine serum albumin (BSA, cat. A2153) were purchased from Sigma-Aldrich. Sulfo-LC-SPDP (cat. 21650), DyLight488-NHS ester (cat. 46402) and DyLight800-NHS ester (cat. 46421) were purchased from Thermo Fisher Scientific. DBCO-PEG-SH (MW 1k, cat. HE048003-1K), SPDP-PEG36-NHS ester (cat. BP-21938) and DBCO-PEG24-NHS ester (cat. BP-25726) were purchased from BroadPharm. Cys(Npys)-TAT-FAM was obtained from AnaSpec (cat. AS-61213). CRYBMIM-Cys(Npys) and functionalized CPP peptides were synthesized by LifeTein LLC, with details listed in supplementary **Table S1**. Amicon Ultra Centrifugal Filters (MWCO 30K, 0.5 ml cat. UFC503096, 15 ml cat. UFC903008) were purchased from MilliporeSigma. C18 solid phase extraction (SPE) cartridge (cat. 60108) and strong cation exchange spin columns (cat. 90009) were purchased from Thermo Fisher Scientific. gRNAs were synthesized and HPLC-purified by IDT Corporation (Newark, NJ, USA).

### Protein expression and purification

Protein purification for 3xNLS-SpCas9 and the V1, V2 and V3 variants were performed similarly to our previous approach (*44*). Briefly, pET21a plasmid backbone for each Cas9 protein was transformed into *E. coli* Rosetta(DE3)pLysS cells (EMD Millipore) for protein production. Following induction with IPTG, cells were pelleted by centrifugation and then resuspended with Nickel-NTA buffer (100 mM Tris(hydroxymethyl)aminomethane (Tris) + 1 M sodium chloride (NaCl) + 20 mM imidazole + 1 mM TCEP, pH 7.5) supplemented with HALT Protease Inhibitor Cocktail, ethylenediaminetetraacetic acid (EDTA)-Free (100×) (ThermoFisher) and lysed with a M-110s Microfluidizer (Microfluidics) following the manufacturer’s instruction. Following purification on Ni-NTA column, protein was eluted using elution buffer (50 mM Tris, 500 mM NaCl, 500 mM imidazole, 10% glycerol, pH 7.5). The Cas9 protein was dialyzed overnight at 4°C in 50 mM 4-(2-Hydroxyethyl)piperazine-1-ethanesulfonic acid (HEPES), 1 M NaCl, 1 mM EDTA, 10% (w/v) glycerol, pH 7.5. Then, the protein was batch bound to loose Capto Q resin (Cytiva) and diluted with 50 mM HEPES, 10% (w/v) glycerol, pH 7.5 until conductivity reached 25 mS/cm. Next, the protein was purified by stacked ion exchange chromatography columns (Columns = a 20ml HiPrep-Q on top of a 5 ml HiTrap-SP, Buffer A = 20 mM HEPES pH 7.5 + 1 mM TCEP, Buffer B = 20 mM HEPES pH 7.5 + 1 M NaCl + 1 mM TCEP, Flow rate = 5 ml/min, CV = column volume = 5 ml). The primary protein peak from the CEC was concentrated in an Ultra-15 Centrifugal Filters Ultracel-30K (Amicon) to a concentration around 100 mM based on absorbance at 280 nm. The purified protein quality was assessed by SDS-PAGE/Coomassie staining to be >95% pure and protein concentration was quantified with Pierce^TM^ BCA Protein Assay Kit (Thermo Fisher Scientific). Protein was stored in –80°C until further use. The sequences of the engineered Cas9 variants are listed in **Table S3**.

### Synthesis of transferrin-DyLight800-TAT-FAM BMC

All reactions were carried out in the reaction buffer containing 20 mM phosphate, 100 mM NaCl, 1 mM EDTA, pH 7.5. The buffer pH was adjusted by using sodium hydroxide (NaOH). All reactions were prepared in 1.5 ml or 5.0 ml capped tubes and rotated end over end at indicated temperatures. The concentrations of the starting materials, intermediates, final products were quantified using NanoDrop 2000 spectrophotometer (Thermo Fisher Scientific). Extinction coefficients are listed in **Tables S1** and **S2**.

The carrier transferrin was first reacted at 125 µM concentration with 2 molar equivalents of DyLight800-NHS ester for 3 h at room temperature. Products were purified by ultrafiltration using 4 x 15 ml reaction buffer with Amicon 30kDa cutoff ultracentrifugal filters to remove excess unreacted DyLight800-NHS ester dye. The resultant Tf-DyLight800 conjugates were then thiolated at 60 µM with 2 molar equivalents of Sulfo-LC-SPDP for 3 h at room temperature. Products were then purified by ultrafiltration with 4 x 15 ml reaction buffer using Amicon 30kDa filters to remove excess unreacted Sulfo-LC-SPDP. The achieved Tf-DyLight800-SPDP was reduced with 5 eq of TCEP at room temperature for 1 h. Products were purified by ultrafiltration with 4 x 15 ml reaction buffer using Amicon 30kDa filters to remove excess TCEP. The resultant Tf-DyLight800-SH was then conjugated at 30 µM with 1.2 eq of Cys(Npys)-TAT-FAM overnight at 4°C.

The resultant Tf-DyLight800-TAT-FAM was purified using a combination of size exclusion chromatography (SEC) and ion exchange chromatography (IEX). Purification was performed on AKTA pure FPLC (Cytiva) at 4°C with UV detection at 280 nm, 495 nm and 700 nm. For SEC purification: column, Superdex 200 Increase 10/300 GL column (Cytiva); flow rate, 0.5 ml/min; mobile phase, 50 mM phosphate, 150 mM NaCl, 10 mM EDTA, pH 8.0; elution, 2 column volume (CV); collection, 1 ml per fraction. The corresponding fractions were collected (determined based on UV absorbance and SDS-PAGE readout) and subjected to IEX purification. For IEX purification: column, MonoS 5/50 GL column (Cytiva); flow rate, 1 ml/min; mobile phase A, 50 mM 3-(*N*-morpholino)propanesulfonic acid (MOPS), 50 mM NaCl, pH 6.5; mobile phase B, 50 mM MOPS, 200 mM NaCl, pH 6.5; method: 5 CV buffer A for loading, 5 CV buffer A for washing, 20 CV 0-100 % buffer B for elution, 5 CV buffer B for cleaning, 5 CV buffer A for equilibration; collection, 0.5 ml per fraction. The corresponding fractions were collected (determined based on UV absorbance and SDS-PAGE readout) and subjected to Amicon 30kDa ultrafiltration for buffer exchange and concentration. 20 mM phosphate, 100 mM NaCl, 1 mM EDTA, pH 7.5 was used as storage buffer.

### Synthesis of transferrin-CRYBMIM BMC

All reactions were carried out in reaction buffer containing 20 mM phosphate, 100 mM NaCl, 1 mM EDTA, pH 7.5, with 10% glycerol. The buffer pH was adjusted by using NaOH. All reactions were set up in 1.5 ml or 5.0 ml capped tubes and rotated end over end at indicated temperature for incubation. The concentrations of the starting materials, intermediates, final products were quantified using NanoDrop 2000 spectrophotometer. Extinction coefficients are listed in **Tables S1** and **S2**.

The cargo CRYBMIM-Cys was freshly prepared by reducing CRYBMIM-Cys(Npys) at 10 mg/ml with 20 mM TCEP followed by solid phase extraction (SPE) purification to remove excess reducing agent. For SPE purification: column, C18 SPE cartridge (Thermo Fisher Scientific); mobile phase A, H_2_O with 0.1% trifluoroacetic acid (TFA); mobile phase B, acetonitrile (ACN) with 0.1% TFA; method, 2 CV 100% B for activation, 3 CV 5% B for equilibration, 3 CV 5% B after loading for removing TCEP, 3 CV 80% B for elution. The collected elution fractions containing CRYBMIM-Cys were dried in a vacuum evaporator (Genevac EZ-2 Elite) and used immediately for conjugation.

The carrier transferrin was first thiolated with sulfo-LC-SPDP at 250 µM a molar ratio of 1:10 for 3 h at room temperature. Ultrafiltration was performed using Amicon 30k filter unit to remove excess unreacted sulfo-LC-SPDP. The achieved Tf-SPDP was then reacted with 4 molar equivalents CRYBMIM-Cys at 45 µM overnight at 4°C. The achieved Tf-CRYBMIM BMC was purified with IEX to get rid of all unreacted Tf-SPDP. For IEX: column, strong cation exchange spin columns (Thermo Fisher Scientific); buffer A, 50 mM MOPS, 50 mM NaCl, pH 6.5, 10% glycerol; buffer B, 50 mM MOPS, 1 M NaCl, pH 6.5, 10% glycerol; method, 10 ml buffer A for equilibration, 3 x 10 ml buffer A after loading for removing unreacted Tf-SPDP, 3 x 10 ml buffer B for elution. The collected eluents were further purified by ultrafiltration with 3 x 15 ml reaction buffer using Amicon 30kDa filters to remove unreacted CRYBMIM-Cys and replaced the buffer to storage buffer 20 mM phosphate, 100 mM NaCl, 1 mM EDTA, pH 7.5 with 10% glycerol.

### Synthesis of Cas9 tripartite BMC

The Cas9 tripartite BMCs were synthesized using bioorthogonal strategies including amine-NHS ester reaction, thiol-maleimide reaction, unsymmetrical disulfide bond formation, and DBCO-azide SPAAC click reaction. All reaction steps were carried out in the reaction buffer containing 50 mM phosphate (sodium phosphate monobasic), 300 mM NaCl, and 50 mM L-arginine hydrochloride. The buffer pH was adjusted to 7.0 using NaOH. All reactions were set up in PCR tubes or 1.5 ml Eppendorf tubes and rotated at 600 rpm on an Eppendorf thermomixer at indicated temperature for incubation. The concentrations of the starting materials, intermediates, final products were quantified using NanoDrop™ 2000 Spectrophotometer. Extinction coefficients are listed in **Tables S1** and **S2**.

For strategy 1 and 2, 50 µM recombinant Cas9 variant was first reacted with tri-functionalized TAT via thiol-maleimide reaction at a molar ratio of 1:12 to prepare the Cas9-TAT conjugates. After 2.5 h of reaction at room temperature, ultrafiltration was performed using Amicon ultra centrifugal filter unit with molecular weight cutoff as 30k (in short as Amicon 30k) to remove excess unreacted TAT. 30 µM KIT antibody (Ab) and 100 µM *apo*-transferrin (Tf), respectively, were reacted with Sulfo-LC-SPDP at a molar ratio of 1:100 to prepare the Ab-/Tf-SPDP. After 1 h of reaction at 37 °C, ultrafiltration was performed using Amicon 30k filter unit to remove excess unreacted Sulfo-LC-SPDP.

For strategy 1, the 50 eq of linker DBCO-PEG-SH was first ligated with 50 µM Cas9-TAT conjugates via SPAAC click reaction. After 3 h of reaction at 37 °C, ultrafiltration was performed using Amicon 30k filter unit to remove excess unreacted DBCO-PEG-SH linker. The achieved Cas9-TAT-SH was further reacted at 30 µM with 1.4 eq of Ab-/Tf-SPDP to form the final Cas9-TAT-Ab or Cas9-TAT-Tf tripartite BMC for 2 h at room temperature.

For strategy 2, 50 eq of linker DBCO-PEG-SH was first ligated with 100 µM Ab-/Tf-SPDP via asymmetrical disulfide bond formation. After 1 h of reaction at 37 °C, ultrafiltration was performed using Amicon 30k filter unit to remove excess unreacted DBCO-PEG-SH linker. Finally, 100 µM Ab-/Tf-DBCO was further reacted with 50 µM Cas9-TAT at the molar ratio of 1.4:1 via SPAAC click reaction to form the final Cas9-TAT-Ab or Cas9-TAT-Tf tripartite BMC for 3 h of reaction at 37 °C.

For strategy 3, 50 µM recombinant Cas9 variant was first reacted with DBCO-PEG24-NHS ester at a molar ratio of 1:20. 30 µM Ab and 100 µM Tf, respectively, were reacted with SPDP-PEG36-NHS ester at a molar ratio of 1:100. Both reactions were incubated for 1 h at 37 °C, and ultrafiltration was performed using Amicon 30k filter unit to remove excess unreacted reagents. 50 µM of the Cas9-PEG-DBCO was then conjugated with the thiolated KIT antibody or transferrin at a ratio of 1:2 to form the Cas9-Ab/Tf dipartite constructs after 1.5 h incubation at 37°C. Finally, 30 µM of the Cas9-Ab/Tf dipartite was reacted with 12 eq of azide-functionalized CPP peptides at 4°C overnight. For CPPs with limited aqueous solubility, such as TAT-Ebola and A5K, peptides were first dissolved in buffer with 5% (w/v) Pluronic F-127, and then mixed with Cas9-Ab/Tf dipartite to confer SPAAC reaction similarly as described above. The Cas9 tripartite BMC was purified with ultrafiltration using Amicon 30k filter unit to remove excess peptides.

### Quantify the CPP Valency of Cas9 Tripartite BMCs

The tri-functionalized TAT used in the synthesis possess a desthiobiotin group at the N-terminus. Valency of desthiobiotin-containing TAT on BMCs could be analyzed using the QuantTag biotin quantitation kit per the manufacturer’s instructions (cat. BDK-2000, Vector Laboratories). Serial dilution of stock solution of desthiobiotin TAT peptides was used as reference standard to establish calibration curves, with the transferrin reaction intermediate (Tf-SPDP) used as the negative control.

### Synthesis of Cas9 tripartite BMC with Dylight488 labeling

DyLight™ 488 NHS ester (cat. 46402, Thermo Fisher Scientific) was dissolved in DMF at 5 mg/ml. To synthesize the control Cas9 protein, 80 µM of the unconjugated Cas9 was labeled with 3 eq of DyLight™ 488 NHS ester and the reaction was incubated for 1 h at 37°C. Similarly, in a separate reaction, 80 µM of unconjugated Cas9 was reacted both with 3 eq of DyLight™ 488 NHS ester and 24 eq of DBCO-PEG24-NHS ester and the reaction was incubated for 1 h at 37°C. The final products were purified by Amicon 30K filter unit to remove excess DBCO-PEG24-NHS and DyLight™ 488. The Cas9 tripartite BMC was further synthesized similarly as mentioned above. The final products, Cas9-DyLight488 and Cas9-Ab-TAT-DyLight488, were confirmed by silver staining and streptavidin staining.

### SDS-PAGE and staining

The reaction products (0.5-1 µg denatured in Laemmli buffer) were analyzed with sodium dodecyl sulfate polyacrylamide gel electrophoresis (SDS-PAGE) using 4-12% polyacrylamide Bis-Tris gels at 130 V for 75 min (Invitrogen). The resolved gels were first washed with 2 × 5 min ultrapure water and stained with Coomassie blue (Bio-Rad, Coomassie brilliant blue R-250 staining solution) or silver stain (Thermo Fisher Scientific, Pierce silver stain kit) per the manufacturer’s instructions. All staining procedures were performed with gentle shaking at room temperature. In short, for Coomassie blue staining, the gel was stained with 15 ml of the staining solution for 1 h, washed with destaining solution (10% acetic acid : 50% methanol : 40% H_2_O) with at least three solvent changes until the background is nearly clear. For silver staining, the gel was first fixed with 2 × 15 min 30% ethanol : 10% acetic acid : 60% H_2_O (15 ml), washed with 2 × 5 min 10% ethanol : 90% H_2_O solution, and subjected to staining per the manufacturer’s instructions. Stained gels were imaged using the Amersham ImageQuant 800 optical digital imaging system (Cytiva).

For analysis of desthiobiotin-containing BMCs, biotinylation was analyzed using SDS-PAGE followed by IRDye 800CW streptavidin staining (1 µg/ml in Intercept blocking buffer (Li-Cor) for 30 min at room temperature. The gels were then washed with Tris buffered saline with Tween-20 (TBST) three times for 5 min and imaged using the Odyssey CLx digital fluorescent imaging system (Li-Cor).

### Mass Photometry

Mass photometry was performed using the TwoMP mass photometer (Refeyn, Oxford, United Kingdom). Calibration was performed using BSA, with all samples analyzed at 250 nM final concentration in the Cas9 reaction buffer (mentioned above) at room temperature. The Refeyn Sample Preparation Pack (6-wells, cat. MP-CON-21008) was used to apply samples onto the coverslip. Carl Zeiss™ Immersol™ 518 F Immersion Oil (SKU: 444960-0000-000) was used on the objective lens. For each sample, 15 µl of sample buffer was first added into the well of cassette and analyzed immediately to find focus and ensure the sharpness is within the appropriate range (5.0-6.5). Then 1 µl of sample was added into the buffer droplet and was mixed thoroughly. Light scattering kinetics were subsequently recorded using Refeyn AcquireMP version 2023 R1.1, and the data were analyzed using Refeyn DiscoverMP version 2023 R1.2 to calculate BMC mass distributions. The yield of Cas9-Tf-TAT BMC was calculated by combining the molecular binding events of the monovalent tripartite (MW=240 kDa) and the multivalent Cas9-Tf-Cas9 construct (MW=400 kDa); similarly, the yield of Cas9-Ab-TAT BMC was calculated by combining the molecular binding events of the tripartite (MW=305 kDa), the multivalent Cas9-Ab-Cas9 construct (MW=465 kDa), and the dimer of the tripartite (MW=610 kDa).

### *In vitro* DNA cleavage assay

Linearized DNA substrate was first prepared by digesting 30 µg of plasmid DNA (final concentration 437 ng/µl) with 300 units of BsaHI restriction enzyme (New England Biolabs) in CutSmart Buffer (New England Biolabs) at 37 °C for 1 hr. The restriction enzyme was heat-inactivated by incubation at 80 °C for 20 minutes. After cooling to room temperature, the linearized plasmid DNA was diluted to 40 nM as working stock using IDTE buffer (10 mM Tris, 0.1 mM EDTA, IDT). RNP complexes were assembled by incubating Cas9 or BMCs stocks (in reaction buffer, 50 mM phosphate, 300 mM NaCl, and 50 mM L-arginine hydrochloride, pH 7.0) with 100 µM purified sgRNA at 1:3 molar ratios for 20 minutes at room temperature. Serially diluted working stock was prepared immediately before use. For stability test, the RNP complex was incubated with reaction buffer (50 mM phosphate, 300 mM NaCl, and 50 mM L-arginine hydrochloride), RPMI+10% FBS, or cell lysates (2 million of Kasumi-1 TLR cells were suspended in PBS and cell lysates were obtained through three freeze-thaw cycles, preserving all protein contents in native forms) at 37 °C for 24 h. Serially diluted working stock was then prepared accordingly.

The cleavage reaction was initiated by mixing linearized plasmid DNA (final concentration 4 nM) with RNP stocks (final concentration ranging from 5 to 80 nM) in nuclease reaction buffer (20 mM HEPES, 100 mM NaCl, 5 mM MgCl_2_, 0.1 mM EDTA, pH 6.5). The reactions were carried out at 37 °C for 30 min, then stopped by addition of 0.4 units of proteinase K (New England Biolabs), followed by incubation at 56 °C for 15 min. Gel electrophoresis was carried out using 1% agarose gels (containing GelRed dye (Biotium), 10 µl in 100 ml gel) in TBE buffer, resolved at 140 V for 75 min. Resolved gels were imaged using Typhoon FLA 9500 fluorescent imaging system (GE Healthcare), with the photomultiplier tube voltage of 750 V, pixel size of 50 µm, and excitation and emission wavelengths of 542 and 568 nm, respectively. Densitometric analysis was performed using ImageJ (Fiji) by quantifying the signal intensities of cleaved and uncleaved DNA bands. The cleavage efficiency for each individual samples was calculated as the percentage of the two cleaved bands over all DNA bands.

### Cell culture

The cell lines MV-411 and Kasumi-1 and HEK293T were obtained from DSMZ and ATCC, respectively. Kasumi-1 TLR and HEK293T TLR cells were generated as described (*51*). The identity of all cell lines was verified by Short Tandem Repeat (STR) analysis (Integrated Genomics Operation Core Facility, MSKCC) and absence of Mycoplasma contamination was determined using MycoAlert mycoplasma detection kit per manufacturer’s instructions (Lonza, cat. LT07-418). MV-411 and Kasumi-1 TLR cells were maintained in RPMI 1640 medium (Corning, cat. 10-040-CV) supplemented with 10% (v/v) fetal bovine serum (FBS, Corning), 1% (v/v) penicillin-streptomycin (100 U/ml penicillin and 100 μg/mL streptomycin, Gibco, cat. 15140122), and 1% (v/v) L-glutamine (4 mM concentration, complete medium). HEK293T TLR cells were maintained in Dulbecco’s Modified Eagle Medium (DMEM, Corning, cat.10-017-CV) supplemented with 10% (v/v) FBS, 1% penicillin-streptomycin, and 1% L-glutamine (4 mM concentration, complete medium). All cells were cultured in a humidified environment with 5% CO_2_ at 37 °C.

### Cell viability analysis

MV-411 cells were resuspended in fresh media and plated at a density of 2×10^5^ cells per well with 95 μl media in 96-well plate. 5 μl of the stock solution for tested samples were added to reach indicated final concentrations. After 5 days of treatment, cell viability was assessed using the CellTiter-Glo Luminescent Viability assay, according to the manufacturer’s instructions (Promega). Luminescence was recorded using the Infinite M1000Pro plate reader with integration time of 250 ms (Tecan).

### Electroporation

The Kasumi-1 TLR cells were washed with PBS and resuspended in buffer R (Invitrogen, cat. MPK10096). 250,000 Kasumi-1 TLR cells were used per condition. RNPs of Cas9 tripartite BMCs were formed by complexing purified Cas9 BMCs and 1.2 eq of sgRNA in the nuclease reaction buffer at the final concentration of 5 µM, and then were mixed with Kasumi-1 TLR cells together with 2 µl buffer R and 1.35 µl of Alt-R electroporation enhancer (IDT, cat. 1075916, final concentration 1.8 µM) to make a final volume of 14.5 µl electroporation mixture. Electroporation tube (Neon) was filled with 3 ml electrolytic buffer E (Invitrogen, cat. MPK10096). 10 μl out of the 14.5 µl electroporation mixture was aspirated into the electroporation tip and electroporated using 10-msec pulses twice at 1600 V using the Neon System (Invitrogen). The electroporated cells were resuspended into antibiotic-free RPMI media and incubated at 37 °C for 3 hours. The cells were then sedimented and resuspended in complete media containing 1% penicillin-streptomycin and incubated in a humidified environment with 5% CO_2_ at 37 °C for 3 days for analysis.

### Intracellular Delivery of Cas9 Tripartite BMC

The sgRNA stock was mixed with electroporation enhancer, 250,000 Kasumi-1 TLR cells, and resuspension buffer R to a final volume of 14.5 µl. The amount of sgRNA corresponds to 2.0 eq of the Cas9 tripartite that would be added later. The final concentration of electroporation enhancer is 1.8 µM. The Neon tube was filled with 3 ml electrolytic buffer E. 10 μl of the sgRNA/cell mixture was uptaken into the Neon pipette tip and inserted into the Neon tube. The electroporation parameters were set as 1600V, 10 ms, 2 pulses. The electroporated cells were resuspended into antibiotic-free RPMI media and incubated at 37 °C for 3 hours. The cells were then resuspended in fresh complete RPMI media containing 1% penicillin-streptomycin, and spiked with corresponding amount of Cas9 tripartite, to a final volume of 20 μl. The tripartite/cell mixture was incubated at 37 °C for 30 min, and the cells were spun down and resuspended into fresh complete RPMI media. The cells were further incubated at 37 °C for 3 days before flow cytometry.

### Flow Cytometry

Cells were sedimented by centrifugation at 400 x g for 5 min and resuspended in ice-cold PBS supplemented with 2.5% (v/v) FBS and 1 mg/ml DAPI (1:10,000 dilution, H3570, Thermo Fisher Scientific), and then transferred into 5 mL cell strainer tubes (Corning). Samples were kept on ice in the dark before test. Cells were analyzed using the LSR Fortessa fluorescence-activated cell scanner (FACS, BD Biosciences), with single fluorophore cells used to define specific compensation settings to minimize FACS channel crosstalk. Cell sorting was performed using the BD FACSymphony™ S6 6-way Cell Sorter (BD Biosciences). FACS data were analyzed using FCS Express version 7.22.0031 (De Novo).

### Confocal Fluorescence Emission Microscopy

Confocal microscopy was performed using Nunc Lab-Tek II 8-well chambered cover glass slides (cat. 155409PK, Thermo Fisher Scientific) or 384-well glass bottom plates (cat. P384-1.5H-N, Cellvis). First, in the case of assessing Tf-Dylight800-TAT-FAM BMC, glass was coated with 0.1% (w/v) solution of poly-L-lysine (Sigma-Aldrich, cat. P8920) for 1 h at room temperature followed with 3 times of PBS washing, or in the case of assessing Cas9 tripartite BMCs, glass was coated with 10 µg/ml fibronectin (cat. F1141-1MG, Sigma Aldrich) in PBS buffer overnight at 4°C. All wells were air-dried at room temperature right before use. MV-411 cells were used for Tf-Dylight800-TAT-FAM BMC. gRNA-preloaded Kasumi-1 TLR cells were used for Cas9 tripartite BMCs (here gRNA duplex was used, other procedure was same as described above). gRNA duplex was formed with crRNA (IDT cat. 306345002) and fluorophore-labeled Alt-R® CRISPR-Cas9 tracrRNA (ATTO™ 647, IDT cat. 10007853) in the duplex buffer (IDT cat. 1072570), annealed at 95 °C for 5 min, and cooled to room temperature.

For Tf-Dylight800-TAT-FAM BMC, MV-411 cells were stained as bulk in complete media supplemented with Hoechst 33342 (final concentration 1 μg/ml, cat. H3570, Thermo Fisher Scientific) and MitoTracker red FM (final concentration 100 nM, cat. M22425, Thermo Fisher Scientific). The stained cells were then distributed to individual wells at a density of 200k cells/300 µl per condition and incubated for 30 min at 37°C for adherence. Tf-Dylight800-TAT-FAM BMC were added at a final concentration of 40 nM and same volume of solvent was used as negative controls.

For Cas9 tripartite BMCs, the gRNA pre-loaded Kasumi-1 TLR cells were first rested in antibiotic-free RPMI media at 37 °C for 3 hours in the chambered slide. Then the Kasumi-1 TLR cells were incubated with DyLight™ 488 labeled Cas9 tripartite BMC and unconjugated Cas9 at a final concentration of 1 µM for 30 min at 37 °C, and stained with Hoechst 33342. The puncta of DyLight488 fluorescence signals per nucleus were quantified via ImageJ (version v1.54p). The confocal image file was first split into individual channels. The threshold of the blue (Hoechst) channel was adjusted and each individual nucleus was recognized as region of interest (ROI). The puncta from the green (DyLight488) channel were recognized as signal maxima with the prominence value set as > 800. The number of puncta was quantified for each nucleus ROI.

SP8 confocal microscope (Leica) equipped with 63X oil objective were used for Tf-Dylight800-TAT-FAM BMC, recording 9 views per condition (tiled scan) with five consecutive z-stacks at 1 µm steps (centered on the widest cellular diameter for the majority of cells). Zeiss LSM 880 Confocal Laser Scanning Microscope (Carl Zeiss AG) with oil objective 63X plus air 20X were used for Cas9 tripartite BMCs, recording n=2 views per condition at single z-stack. Time lapse frames were taken at 10 min per frame for a duration of 4 hours. The images were analyzed using ImageJ-FIJI and ZEN Lite.

### DNA sequencing and TIDE analysis

Genomic DNA was extracted from 400k sorted cells using Invitrogen PureLink Genomic DNA Mini Kit per manufacturer’s instructions (cat. K182002, Thermo Fisher Scientific). The elution was repeated twice using 25 μl elution buffer. DNA concentration was measured using the NanoDrop 2000 spectrophotometer, and diluted to 50 ng/μl with nuclease-free water. PCR was performed using the Vapo.Protect Mastercycler Pro thermal cycler (Eppendorf) with 50 ng template DNA, 1 μM forward and reverse primers, 0.2 µM NTPs (cat. 18427013, Invitrogen), and 1.25 units of Ex Taq DNA Polymerase (Takara Bio Inc.). PCR products was resolved using 1% agarose gel electrophoresis, and the product DNA bands were extracted using Invitrogen PureLink Quick Gel Extraction kit per the manufacturer’s instructions (cat. K210012, Thermo Fisher Scientific). Purified DNA was sequenced using the Sanger method (Eton Bioscience Inc.), and sequencing data were analyzed using the Tracking of Indels by DEcomposition (TIDE) method(*52*).

### Statistical analysis

Numerical and statistical analyses were performed using OriginPro (OriginLab, Northampton, MA). Statistical significance was determined using two-tailed t-tests for continuous pair variables, and ANOVA with Bonferroni and Tukey’s test for multiple comparisons.

## Acknowledgements

We thank all our laboratory members for helpful suggestions and critical comments on the manuscript, Zhi Zheng, Aidin Shabro, Eric Chan, Eric Rosiek, Fang Fang, Gabriel Barretto, Magdalena Parys, Rong Wang, and George Sukenick, Chunyan Ren, Sabine Studer for technical assistance, and Jason Lewis for comments on the manuscript. HMM is supported by the Aubrey Fund for Pediatric Cancer Research and the Tow Center for Developmental Oncology. AK is a Scholar of the Leukemia & Lymphoma Society and acknowledges generous support of multiple funders listed below.

## Author Contributions

Conceptualization: AK, DL, NW, SAW, DEB

Methodology: AK, DL, NW, HMF, JR, SR, SAM, KP, MY

Investigation: All authors Visualization: AK, DL, NW

Funding acquisition: AK, SAW, DEB

Project administration: AK, SAW, DEB

Supervision: AK, SAW, DEB

Writing – original draft: AK, DL, NW; final draft, all authors

## Data and materials availability

All materials generated in this study are available upon request subject to a standard Material Transfer Agreement (MTA) provided by MSK.

## Supplementary Materials

The PDF file includes:

Figs. S1 to S17

Tables S1 to S6

Other Supplementary Material for this manuscript includes the following: Movie S1

## References

1. G. Walsh, E. Walsh, Biopharmaceutical benchmarks 2022. Nat Biotechnol 40, 1722–1760 (2022).

2. B. Leader, Q. J. Baca, D. E. Golan, Protein therapeutics: a summary and pharmacological classification. Nat Rev Drug Discov 7, 21–39 (2008).

3. S. B. Ebrahimi, D. Samanta, Engineering protein-based therapeutics through structural and chemical design. Nat Commun 14, 2411 (2023).

4. J. R. Kintzing, M. V. Filsinger Interrante, J. R. Cochran, Emerging strategies for developing next-generation protein therapeutics for cancer treatment. Trends Pharmacol Sci 37, 993–1008 (2016).

5. H. Nada, Y. Choi, S. Kim, K. S. Jeong, N. A. Meanwell, K. Lee, New insights into protein–protein interaction modulators in drug discovery and therapeutic advance. Signal Transduct Target Ther 9, 341 (2024).

6. X. Xie, T. Yu, X. Li, N. Zhang, L. J. Foster, C. Peng, W. Huang, G. He, Recent advances in targeting the “undruggable” proteins: from drug discovery to clinical trials. Signal Transduct Target Ther 8 (2023).

7. A. Kakkar, G. Traverso, O. C. Farokhzad, R. Weissleder, R. Langer, Evolution of macromolecular complexity in drug delivery systems. Nat Rev Chem 1, Article 0063 (2017).

8. J. Yang, J. Kopecek, Macromolecular therapeutics. Journal of controlled release 190, 288–303 (2014).

9. T. Rath, K. Baker, J. A. Dumont, R. T. Peters, H. Jiang, S. W. Qiao, W. I. Lencer, G. F. Pierce, R. S. Blumberg, Fc-fusion proteins and FcRn: Structural insights for longer-lasting and more effective therapeutics. Crit Rev Biotechnol 35, 235–254 (2015).

10. Y. Fu, R. Tang, X. Zhao, Engineering cytokines for cancer immunotherapy: a systematic review. Front Immunol 14, 1218082 (2023).

11. A. Chan, A. Tsourkas, Intracellular protein delivery: approaches, challenges, and clinical applications. BME Front 5, 0035 (2024).

12. A. Boufridi, R. J. Quinn, Harnessing the Properties of Natural Products. Annu Rev Pharmacol Toxicol 58, 451–470 (2018).

13. S. Yan, J. Na, X. Liu, P. Wu, Different targeting ligands-mediated drug delivery systems for tumor therapy. Pharmaceutics 16, 248 (2024).

14. M. Asher, Maturing antibody-drug conjugate pipeline hits 30. Nat Rev Drug Discov 12, pages329–332 (2013).

15. M. J. Birrer, K. N. Moore, I. Betella, R. C. Bates, Antibody-drug conjugate-based therapeutics : state of the science key elements of ADC design. JNCI J Natl Cancer Inst 111, 538–549 (2019).

16. C. Dumontet, J. M. Reichert, P. D. Senter, J. M. Lambert, A. Beck, Antibody–drug conjugates come of age in oncology. Nat Rev Drug Discov 22, 641–661 (2023).

17. I. Porello, F. Cellesi, Intracellular delivery of therapeutic proteins. New advancements and future directions. Front Bioeng Biotechnol 11, 1211798 (2023).

18. M. Lindgren, M. Hällbrink, A. Prochiantz, Ü. Langel, Cell-penetrating peptides. Trends Pharmacol Sci 21, 99–103 (2000).

19. P. A. Wender, D. J. Mitchell, K. Pattabiraman, E. T. Pelkey, L. Steinman, J. B. Rothbard, The design, synthesis, and evaluation of molecules that enable or enhance cellular uptake: Peptoid molecular transporters. PNAS 97, 13003–13008 (2000).

20. M. Green, P. M. Loewenstein, Autonomous functional domains of chemically synthesized human immunodeficiency virus tat trans-activator protein. Cell 55, 1179–1188 (1988).

21. A. D. Frankel, C. O. Pabo, Cellular uptake of the tat protein from human immunodeficiency virus. Cell 55, 1189–1193 (1988).

22. D. Derossi, A. H. Joliot, G. Chassaing, A. Prochiantz, The third helix of the Antennapedia homeodornain translocates through biological membranes. 269, 10444–10450 (1994).

23. E. Vives, P. Brodin, B. Lebleu, A truncated HIV-1 Tat protein basic domain rapidly translocates through the plasma membrane and accumulates in the cell nucleus. J Biol Chem 272, 16010–16017 (1997).

24. N. Wang, N. A. Mcneer, E. Eton, J. Fass, A. Kentsis, Proteomic barcoding patform for macromolecular screening and delivery. J Proteome Res 23, 2067–2077 (2024).

25. S. Reissmann, M. P. Filatova, New generation of cell-penetrating peptides: Functionality and potential clinical application. Journal of Peptide Science 27, e3300 (2021).

26. J. D. Hirsch, L. Eslamizar, B. J. Filanoski, N. Malekzadeh, R. P. Haugland, J. M. Beechem, R. P. Haugland, Easily reversible desthiobiotin binding to streptavidin, avidin, and other biotin-binding proteins: uses for protein labeling, detection, and isolation. Anal Biochem 308, 343–357 (2002).

27. S. Zhang, J. Shen, D. Li, Y. Cheng, Strategies in the delivery of Cas9 ribonucleoprotein for CRISPR/Cas9 genome editing. Theranostics 11, 614–648 (2020).

28. C. Liu, L. Zhang, H. Liu, K. Cheng, Delivery strategies of the CRISPR-Cas9 gene-editing system for therapeutic applications. Journal of Controlled Release 266, 17– 26 (2017).

29. T. Li, Y. Yang, H. Qi, W. Cui, L. Zhang, X. Fu, X. He, M. Liu, P. feng Li, T. Yu, CRISPR/Cas9 therapeutics: progress and prospects. Signal Transduct Target Ther 8, 36 (2023).

30. M. Pacesa, L. Loeff, I. Querques, L. M. Muckenfuss, M. Sawicka, M. Jinek, R-loop formation and conformational activation mechanisms of Cas9. Nature 609, 191– 196 (2022).

31. H. Nishimasu, F. A. Ran, P. D. Hsu, S. Konermann, S. I. Shehata, N. Dohmae, R. Ishitani, F. Zhang, O. Nureki, Crystal structure of Cas9 in complex with guide RNA and target DNA. Cell 156, 935–949 (2014).

32. S. Yang, S. H. Im, J. Y. Chung, J. Lee, K. H. Lee, Y. K. Kang, H. J. Chung, An antibody-CRISPR/Cas conjugate platform for target-specific delivery and gene editing in cancer. Advanced Science 11, 202308763 (2024).

33. K. Chen, E. C. Stahl, M. H. Kang, B. Xu, R. Allen, M. Trinidad, J. A. Doudna, Engineering self-deliverable ribonucleoproteins for genome editing in the brain. Nat Commun 15 (2024).

34. I. Petta, S. Lievens, C. Libert, J. Tavernier, K. De Bosscher, Modulation of protein– protein interactions for the development of novel therapeutics. Molecular Therapy 24, 707–718 (2015).

35. A. E. Modell, S. L. Blosser, P. S. Arora, Systematic targeting of protein–protein interactions. Trends Pharmacol Sci 37, 702–713 (2016).

36. D. P. Ryan, J. M. Matthews, Protein–protein interactions in human disease. Curr Opin Struct Biol 15, 441–446 (2005).

37. K. Ramaswamy, L. Forbes, G. Minuesa, T. Gindin, F. Brown, M. G. Kharas, A. V Krivtsov, S. A. Armstrong, E. Still, E. De Stanchina, B. Knoechel, R. Koche, A. Kentsis, Peptidomimetic blockade of MYB in acute myeloid leukemia. Nat Commun 9, 110 (2018).

38. S. Takao, L. Forbes, M. Uni, S. Cheng, J. Mario, B. Pineda, Y. Tarumoto, P. Cifani, G. Minuesa, C. Chen, M. G. Kharas, R. K. Bradley, C. R. Vakoc, R. P. Koche, A. Kentsis, Convergent organization of aberrant MYB complex controls oncogenic gene expression in acute myeloid leukemia. Elife 10, e65905 (2021).

39. L. Breda, T. E. Papp, M. P. Triebwasser, A. Yadegari, M. T. Fedorky, N. Tanaka, O. Abdulmalik, G. Pavani, Y. Wang, S. A. Grupp, S. T. Chou, H. Ni, B. L. Mui, Y. K. Tam, D. Weissman, S. Rivella, H. Parhiz, In vivo hematopoietic stem cell modification by mRNA delivery. Science (1979) 381, 436–443 (2023).

40. D. Shi, S. Toyonaga, D. G. Anderson, In vivo RNA delivery to hematopoietic stem and progenitor cells via targeted lipid nanoparticles. Nano Lett 23, 2938–2944 (2023).

41. L. J. Cruz, S. Rezaei, F. Grosveld, S. Philipsen, C. Eich, Nanoparticles targeting hematopoietic stem and progenitor cells: Multimodal carriers for the treatment of hematological diseases. Front Genome Ed 4, 1030285 (2022).

42. D. V. Foss, J. J. Muldoon, D. N. Nguyen, D. Carr, S. U. Sahu, J. M. Hunsinger, S. K. Wyman, N. Krishnappa, R. Mendonsa, E. V. Schanzer, B. R. Shy, V. S. Vykunta, V. Allain, Z. Li, A. Marson, J. Eyquem, R. C. Wilson, Peptide-mediated delivery of CRISPR enzymes for the efficient editing of primary human lymphocytes. Nat Biomed Eng 7, 647–660 (2023).

43. C. Cottet-Rousselle, X. Ronot, X. Leverve, J. F. Mayol, Cytometric assessment of mitochondria using fluorescent probes. Cytometry Part A 79, 405–425 (2011).

44. Y. Wu, J. Zeng, B. P. Roscoe, P. Liu, Q. Yao, C. R. Lazzarotto, K. Clement, M. A. Cole, K. Luk, C. Baricordi, A. H. Shen, C. Ren, E. B. Esrick, J. P. Manis, D. M. Dorfman, D. A. Williams, A. Biffi, C. Brugnara, L. Biasco, C. Brendel, L. Pinello, S. Q. Tsai, S. A. Wolfe, D. E. Bauer, Highly efficient therapeutic gene editing of human hematopoietic stem cells. Nat Med 25, 776–783 (2019).

45. J. S. Chen, Y. S. Dagdas, B. P. Kleinstiver, M. M. Welch, A. A. Sousa, L. B. Harrington, S. H. Sternberg, J. K. Joung, A. Yildiz, J. A. Doudna, Enhanced proofreading governs CRISPR-Cas9 targeting accuracy. Nature 550, 407–410 (2017).

46. S. H. Sternberg, B. Lafrance, M. Kaplan, J. A. Doudna, Conformational control of DNA target cleavage by CRISPR-Cas9. Nature 527, 110–113 (2015).

47. M. Jinek, K. Chylinski, I. Fonfara, M. Hauer, J. A. Doudna, E. Charpentier, A programmable dual-RNA-guided DNA endonuclease in adaptive bacterial immunity. Science (1979) 337, 816–821 (2012).

48. F. Jiang, D. W. Taylor, J. S. Chen, J. E. Kornfeld, K. Zhou, A. J. Thompson, E. Nogales, J. A. Doudna, Structures of a CRISPR-Cas9 R-loop complex primed for DNA cleavage. Science (1979) 351, 863–867 (2016).

49. A. Dhabaria, P. Cifani, C. Reed, H. Steen, A. Kentsis, A high-efficiency cellular extraction system for biological proteomics. J Proteome Res 14, 3403–3408 (2015).

50. M. T. Certo, B. Y. Ryu, J. E. Annis, M. Garibov, J. Jarjour, D. J. Rawlings, A. M. Scharenberg, Tracking genome engineering outcome at individual DNA breakpoints. Nat Methods 8, 671–676 (2011).

51. S. Iyer, A. Mir, J. Vega-Badillo, B. P. Roscoe, R. Ibraheim, L. J. Zhu, J. Lee, P. Liu, K. Luk, E. Mintzer, D. Guo, J. Soares De Brito, C. P. Emerson, P. D. Zamore, E. J. Sontheimer, S. A. Wolfe, Efficient homology-directed repair with circular single-stranded DNA donors. CRISPR J 5, 685–701 (2022).

52. E. K. Brinkman, T. Chen, M. Amendola, B. Van Steensel, Easy quantitative assessment of genome editing by sequence trace decomposition. Nucleic Acids Res 42 (2014).

