## Supplementary materials for "Modular platform for therapeutic drug delivery using trifunctional bio-orthogonal macromolecular conjugates"

**This PDF file includes:**

Figs. S1 to S17

Tables S1 to S6

**Other Supplementary Materials for this manuscript include the following:**

Movie S1

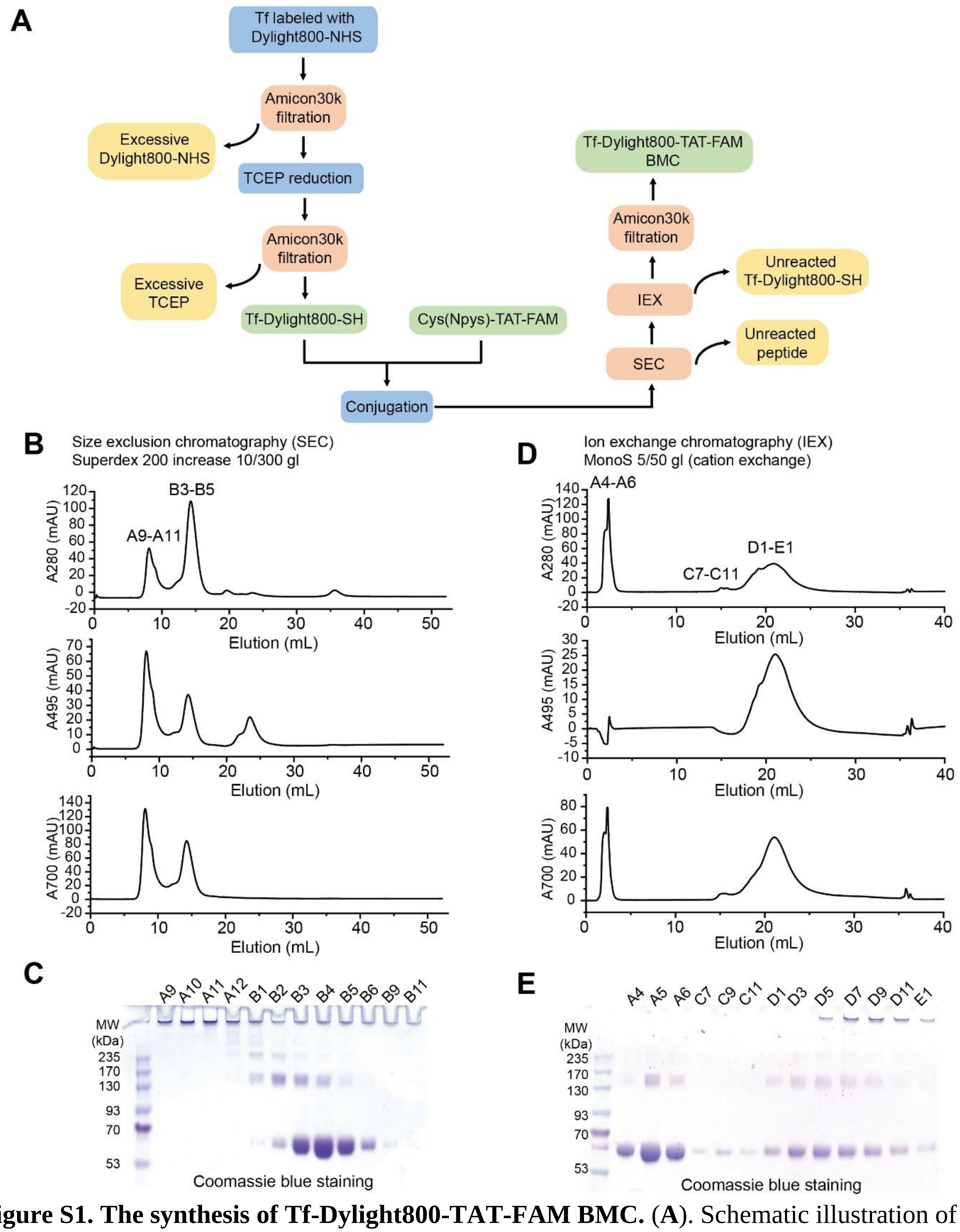


**Figure S1. The synthesis of Tf-Dylight800-TAT-FAM BMC.** (**A**). Schematic illustration of the synthetic strategy and downstream processing procedures to construct Tf-Dylight800-TAT-FAM BMC. (**B**). Size exclusion chromatography purification of Tf-Dylight800-TAT-FAM BMC. Absorbance at 280 nm, 495 nm and 700 nm were used to monitor the elution of Tf-Dylight800-TAT-FAM BMC. (**C**). The corresponding SDS-PAGE gel with Coomassie Blue staining monitored the elution of Tf-Dylight800-TAT-FAM BMC. Fractions B3-B5 were collected for downstream analysis. (**D**). Cation exchange chromatography purification of Tf-Dylight800-TAT-FAM. Absorbance at 280 nm, 495 nm and 700 nm were used to monitor the elution of Tf-Dylight800-TAT-FAM BMC. (**E**). The corresponding SDS-PAGE gel with Coomassie Blue staining monitored the elution of Tf-Dylight800-TAT-FAM BMC. Fractions D1-E1 were collected as final product for biological studies.

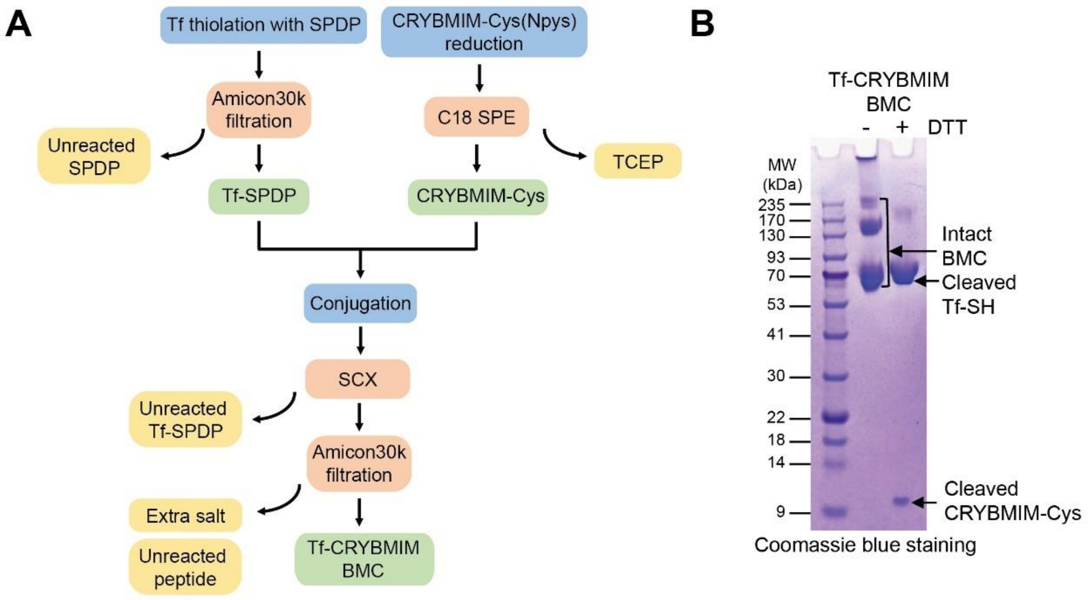


**Figure S2. The synthesis of Tf-CRYBMIM BMC.** (**A**). Schematic illustration of the synthetic strategy and downstream processing procedures to construct Tf-CRYBMIM BMC. (**B**). The final product Tf-CRYBMIM BMC was assessed using SDS-PAGE gel with Coomassie Blue staining. Addition of DTT (final concentration 100 mM) reduced the disulfide bond linker and cleaved off the cargo CRYBMIM, demonstrating successful conjugation of the CRYBMIM onto transferrin carrier to form the Tf-CRYBMIM BMC.

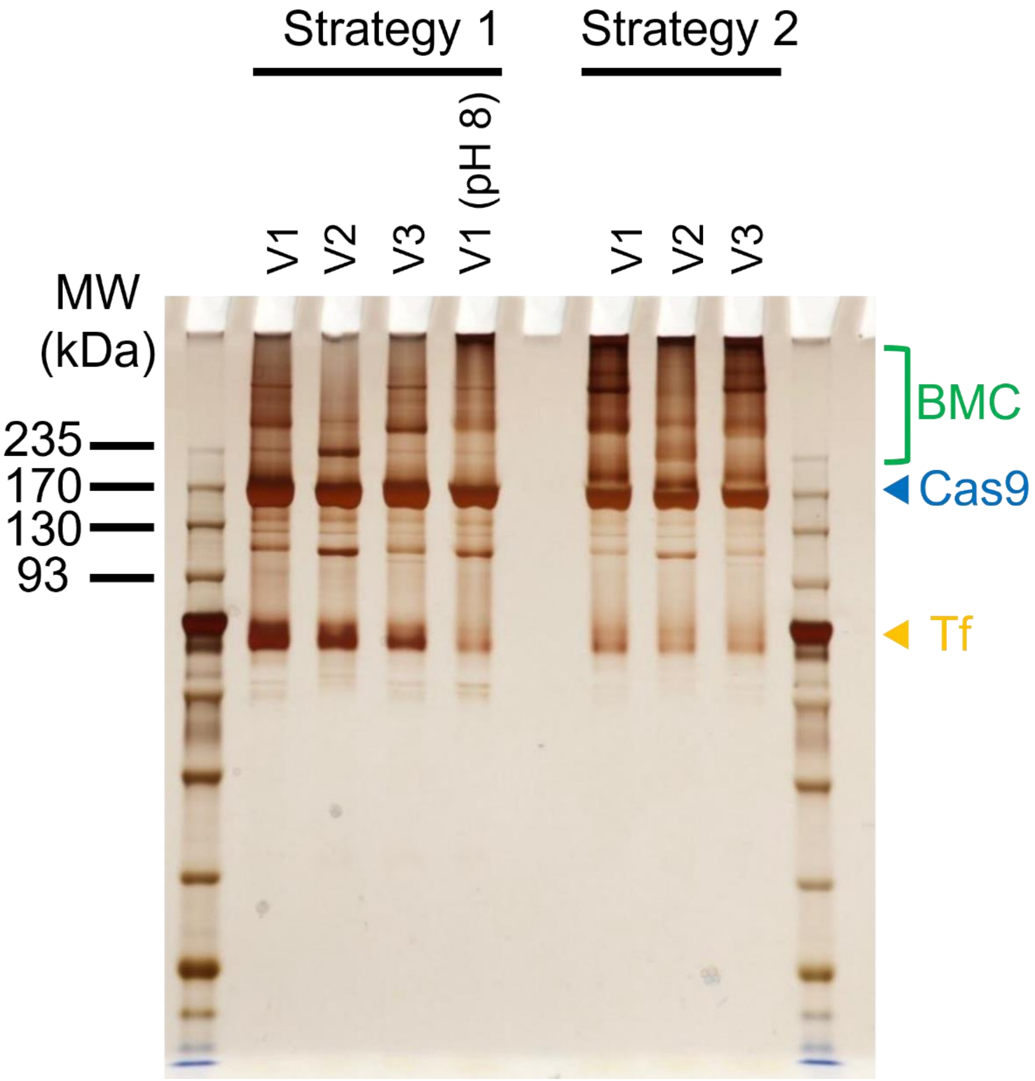


**Figure S3. Cas9 tripartite BMCs synthesized by Strategies 1 and 2 with moderate yields**. Silver staining of the SDS-PAGE showed the reaction intermediates and final products with three SpCas9 variants (V1, V2 & V3) that differ in the position of the cysteine within the polypeptide. pH =8.0 was tested for Cas9 V1 to investigate the influence of pH on reaction yields. The reaction pH was kept at 7.0 for all the other conditions.

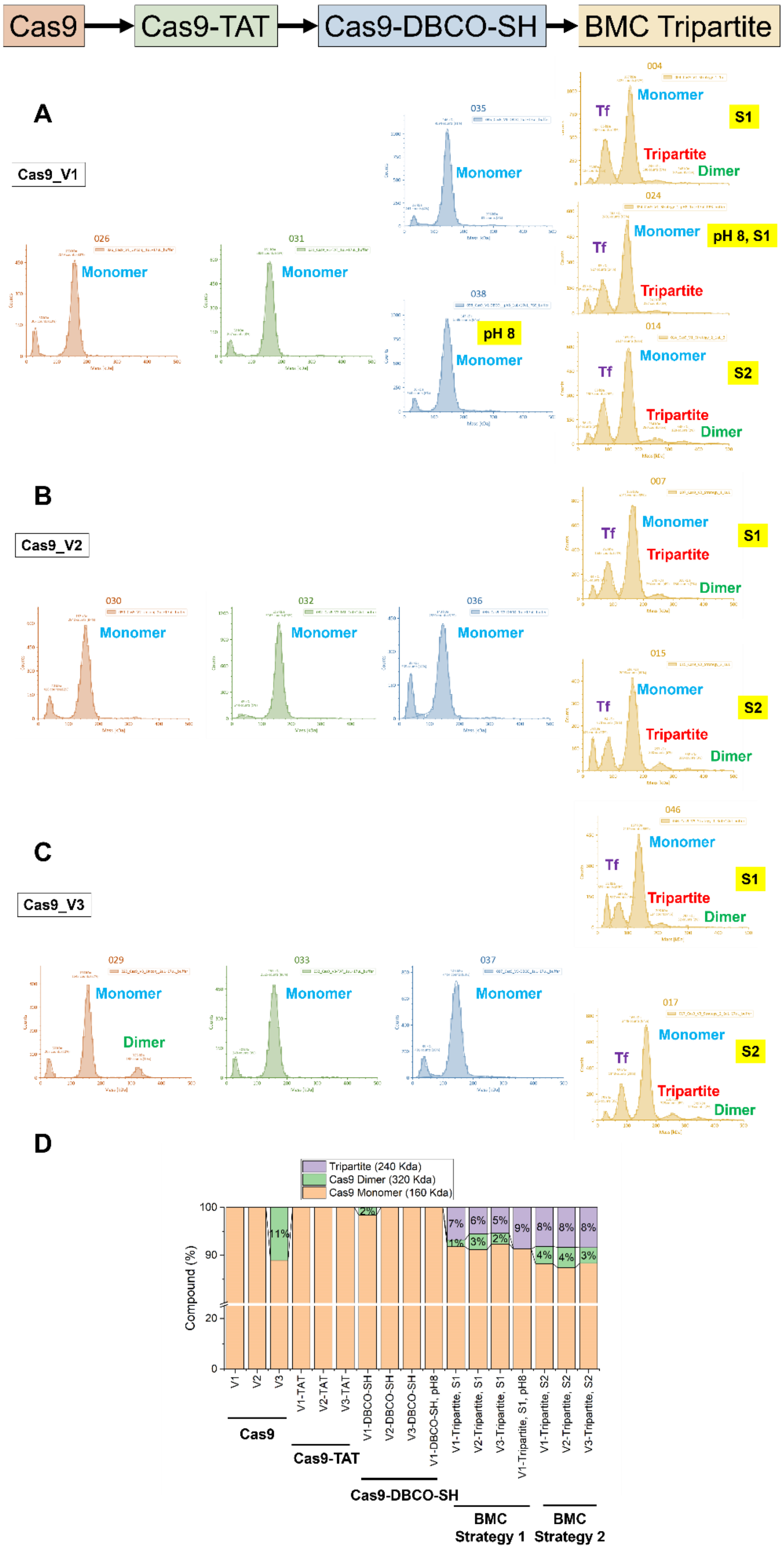

Cas9
Cas9-TAT
Cas9-DBCO-SH
BMC Tripartite
A
Cas9_V1
026
031
035
038
004
024
014
Monomer
pH 8
Tf
Tripartite
Dimer
S1
pH 8, S1
S2
B
Cas9_V2
030
032
036
007
015
C
Cas9_V3
029
033
037
046
017
D
Tripartite (240 Kda)
Cas9 Dimer (320 Kda)
Cas9 Monomer (160 Kda)
Compound (%)
100
90
20
0
11%
2%
7%
1%
6%
3%
5%
2%
9%
8%
4%
8%
4%
8%
3%
V1
V2
V3
V1-TAT
V2-TAT
V3-TAT
V1-DBCO-SH
V2-DBCO-SH
V3-DBCO-SH
V1-DBCO-SH, pH8
V1-Tripartite, S1
V2-Tripartite, S1
V3-Tripartite, S1
V1-Tripartite, S1, pH8
V1-Tripartite, S2
V2-Tripartite, S2
V3-Tripartite, S2
Cas9
Cas9-TAT
Cas9-DBCO-SH
BMC Strategy 1
BMC Strategy 2

**Figure S4. Mass photometry confirmation of Cas9 tripartite BMCs synthesized by Strategies 1 and 2.** (**A−C**) Mass photometry of the reaction intermediates of Cas9 variants V1, V2, and V3. (**D**) Quantitative analysis of the mass photometry data in A−C. All the samples were measured at 250 nM concentration, showing relative abundance as a function of measured molecular mass.

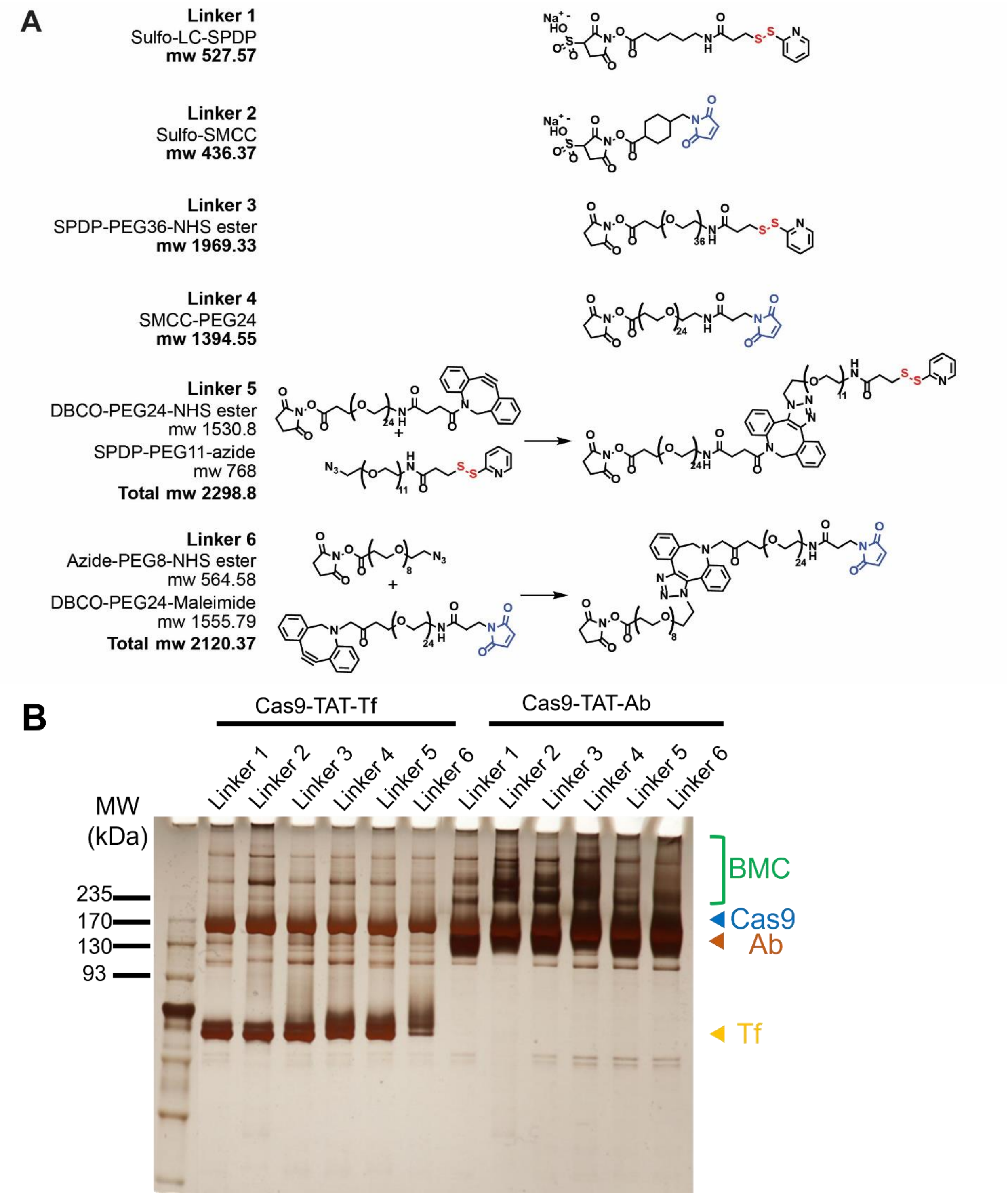


**Figure S5. A variety of chemical linkers were tested to investigate the influence of linker length and reaction reversibility.** (**A**) The molecular structures of all the linkers tested. (**B**) Silver staining of the SDS-PAGE showed that some of the linkers, such as linkers 2-4, improved the BMC yields, albeit moderately.

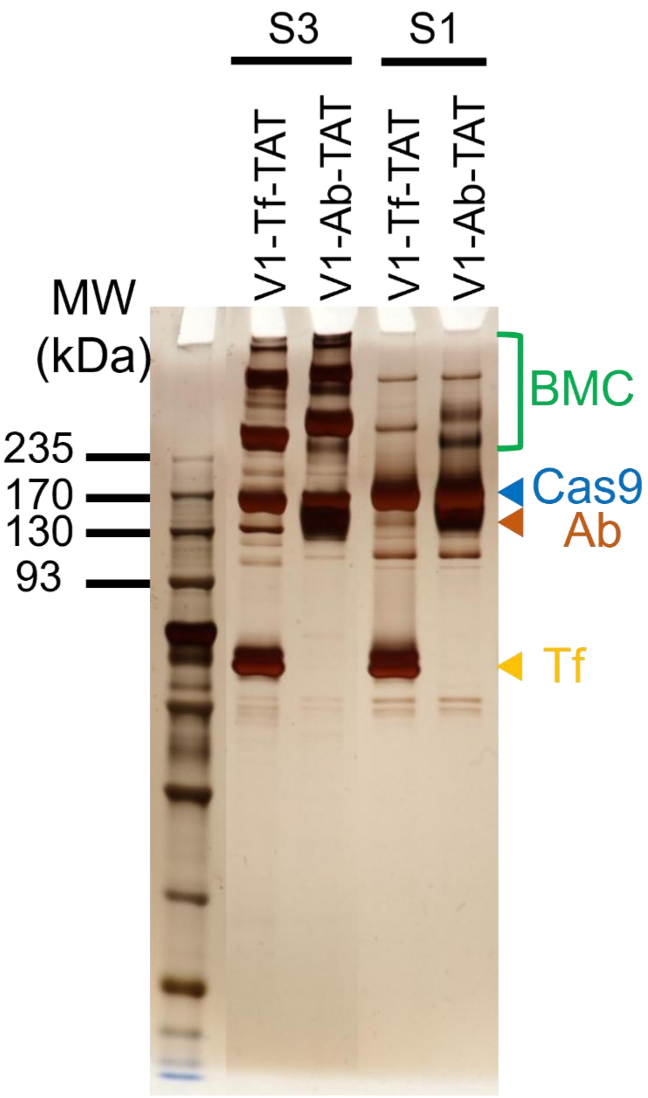


**Figure S6. Strategy 3 improves the yield of Cas9 tripartite BMCs as compared to strategy 1.** S1: strategy 1; S3: strategy 3.

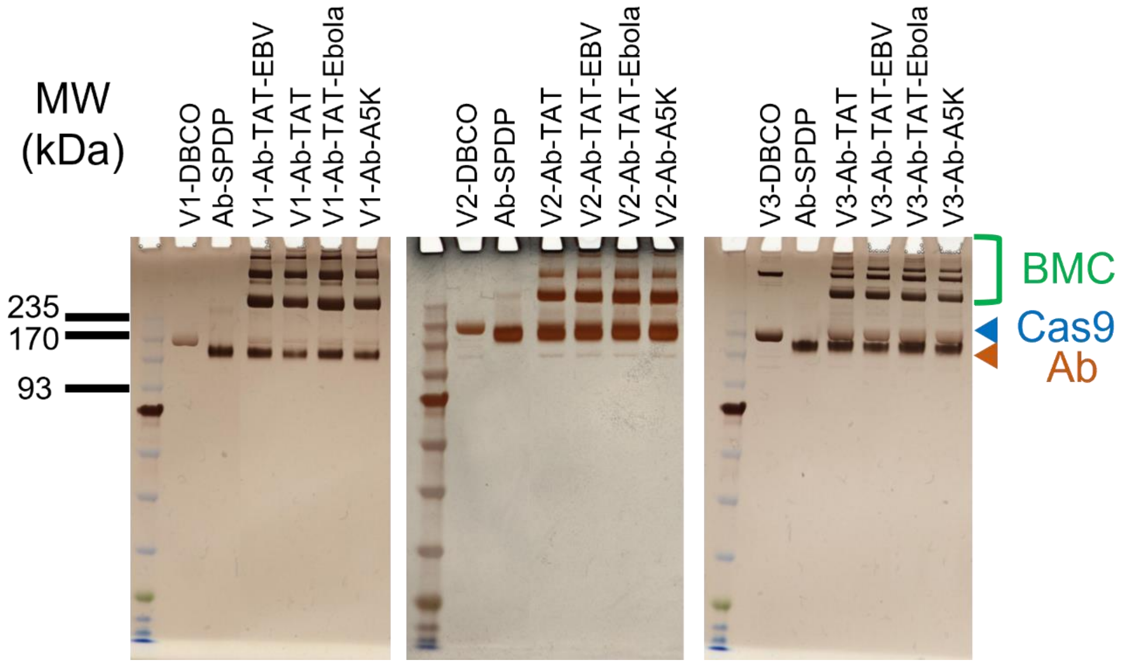


**Figure S7. Cas9 BMCs were synthesized with high yields for all three Cas9 variants and across all the CPPs tested in Strategy 3.**

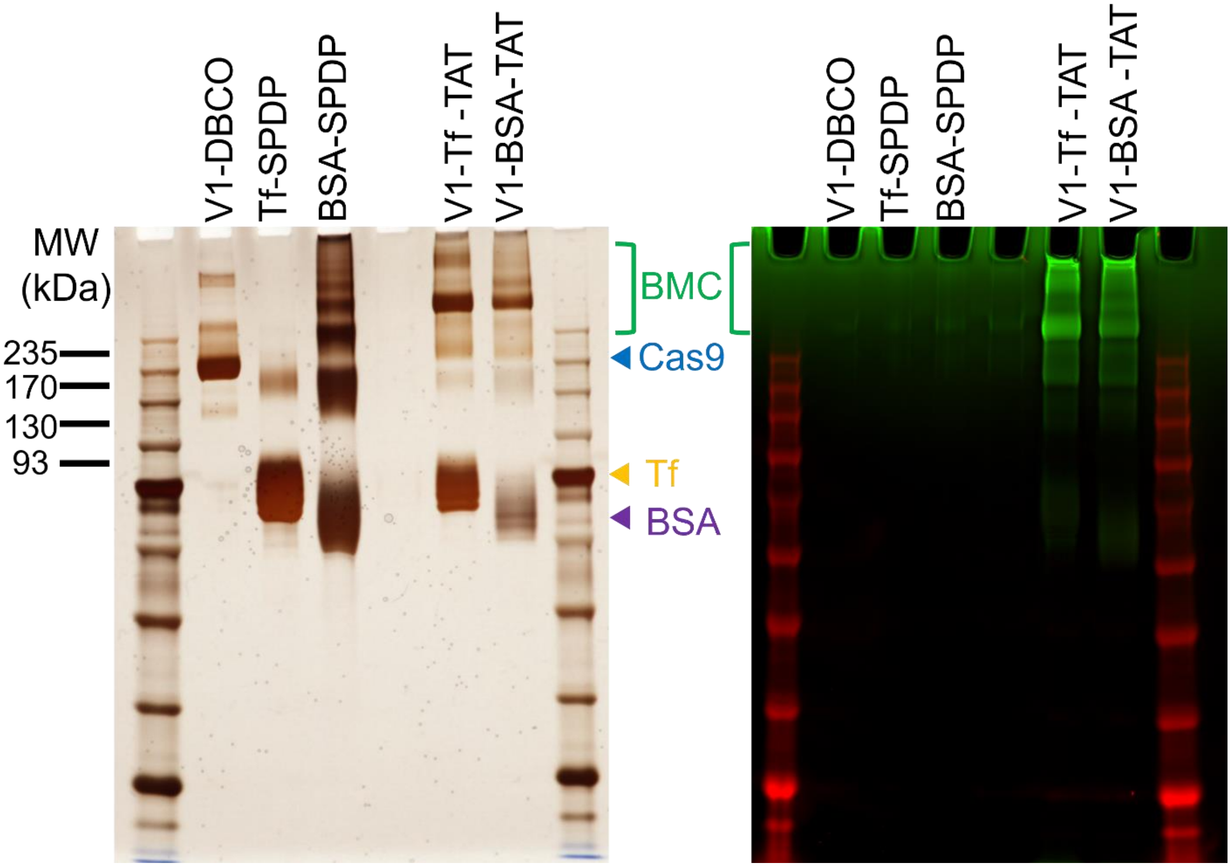


**Figure S8. The developed modular strategy is permissive for diverse carriers.** The silver staining and streptavidin staining confirmed that the Cas9-BSA-TAT BMC was synthesized with high yield and CPP modification.

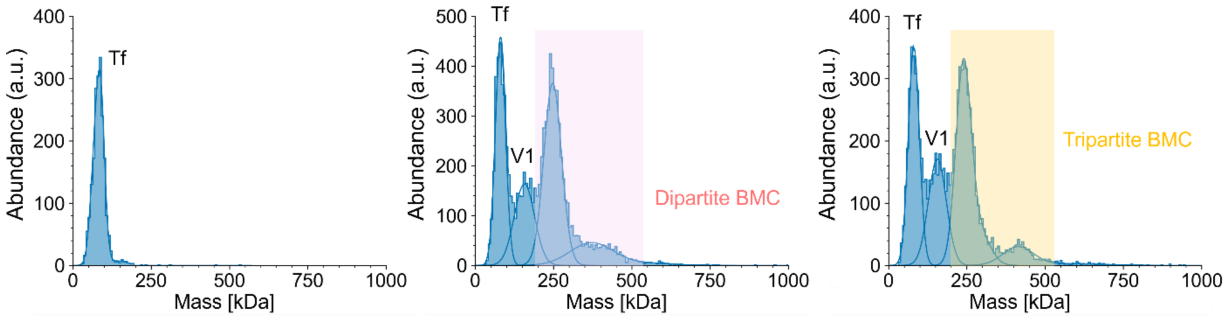


**Figure S9. Mass photometry histograms of the reaction intermediates and final product of the Cas9-Tf-TAT BMC.** The concentration of each sample is 250 nM based on either Cas9 or transferrin. In the final product solution, the expected tripartite (MW=240 kDa) as well as a Cas9-Tf-Cas9 oligomer (MW=400 kDa) were observed.

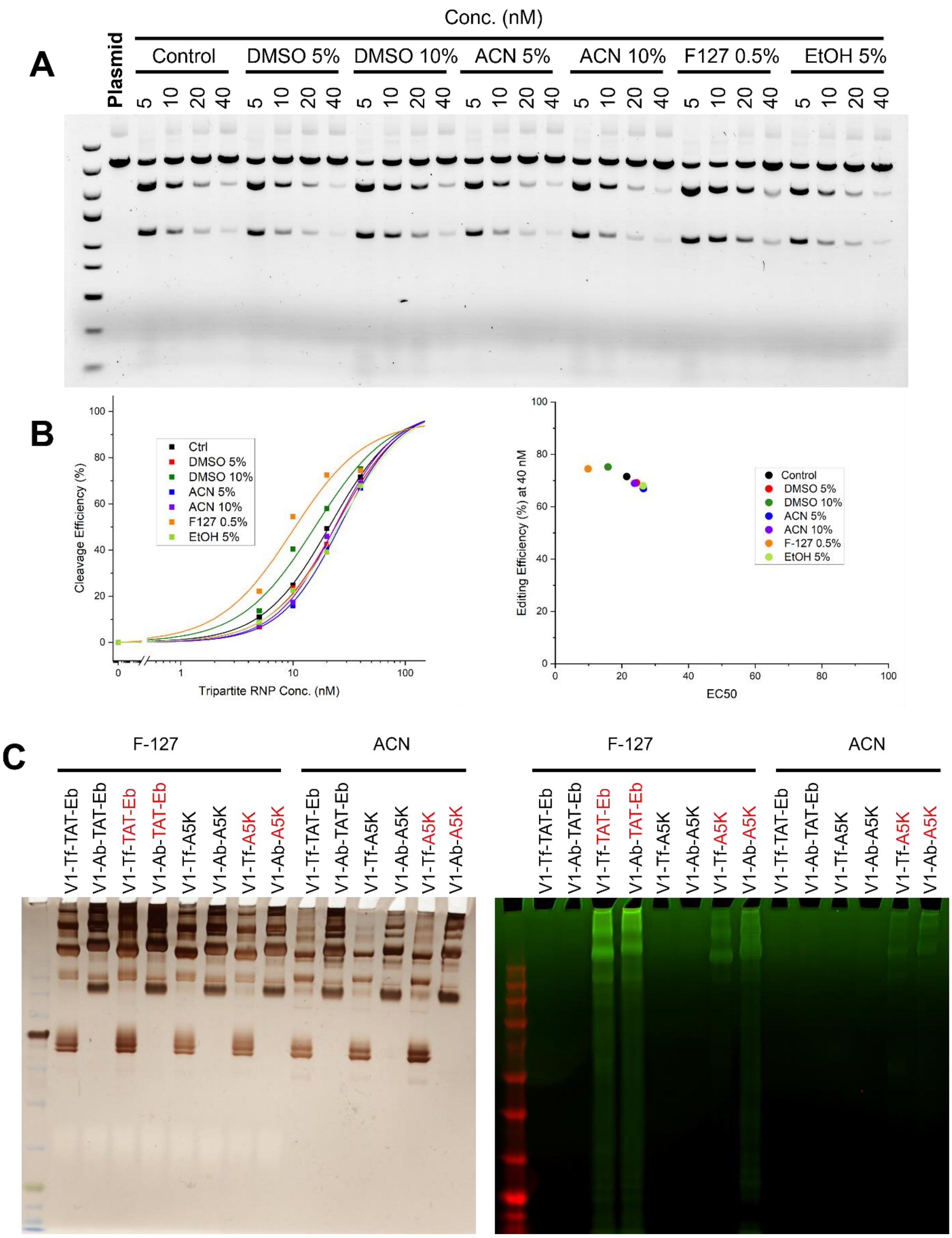
A
Conc. (nM)
Plasmid
Control
DMSO 5%
DMSO 10%
ACN 5%
ACN 10%
F127 0.5%
EtOH 5%
5 10 20 40
B
Cleavage Efficiency (%)
Tripartite RNP Conc. (nM)
Ctrl
DMSO 5%
DMSO 10%
ACN 5%
ACN 10%
F127 0.5%
EtOH 5%
Editing Efficiency (%) at 40 nM
EC50
Control
DMSO 5%
DMSO 10%
ACN 5%
ACN 10%
F-127 0.5%
EtOH 5%
C
F-127
ACN
V1-Tf-TAT-Eb
V1-Ab-TAT-Eb
V1-Tf-TAT-Eb
V1-Ab-TAT-Eb
V1-Tf-A5K
V1-Ab-A5K
V1-Tf-A5K
V1-Ab-A5K

**Figure S10. Optimization on the reaction solvents enhanced the solubility of hydrophobic CPPs and reaction yields.** (**A**) Organic solvents, including DMSO (dimethyl sulfoxide), ACN (acetonitrile) and EtOH (ethanol), as well as the amphipathic block copolymer Pluronic F-127, were tested *in vitro* via plasmid cleavage assay to evaluate the influence of these solvents on the activity of Cas9. (**B**) The plasmid cleavage assay confirmed that overnight incubation at 4°C with these solvents preserves the nuclease activity of Cas9. (**C**) These solvents were then tested on improving the solubility of various CPPs and synthesizing Cas9 tripartite BMCs. Based on the results, Pluronic F-127 showed the best solubilization of CPPs and was therefore used in the synthesis of tripartite Cas9 BMCs.

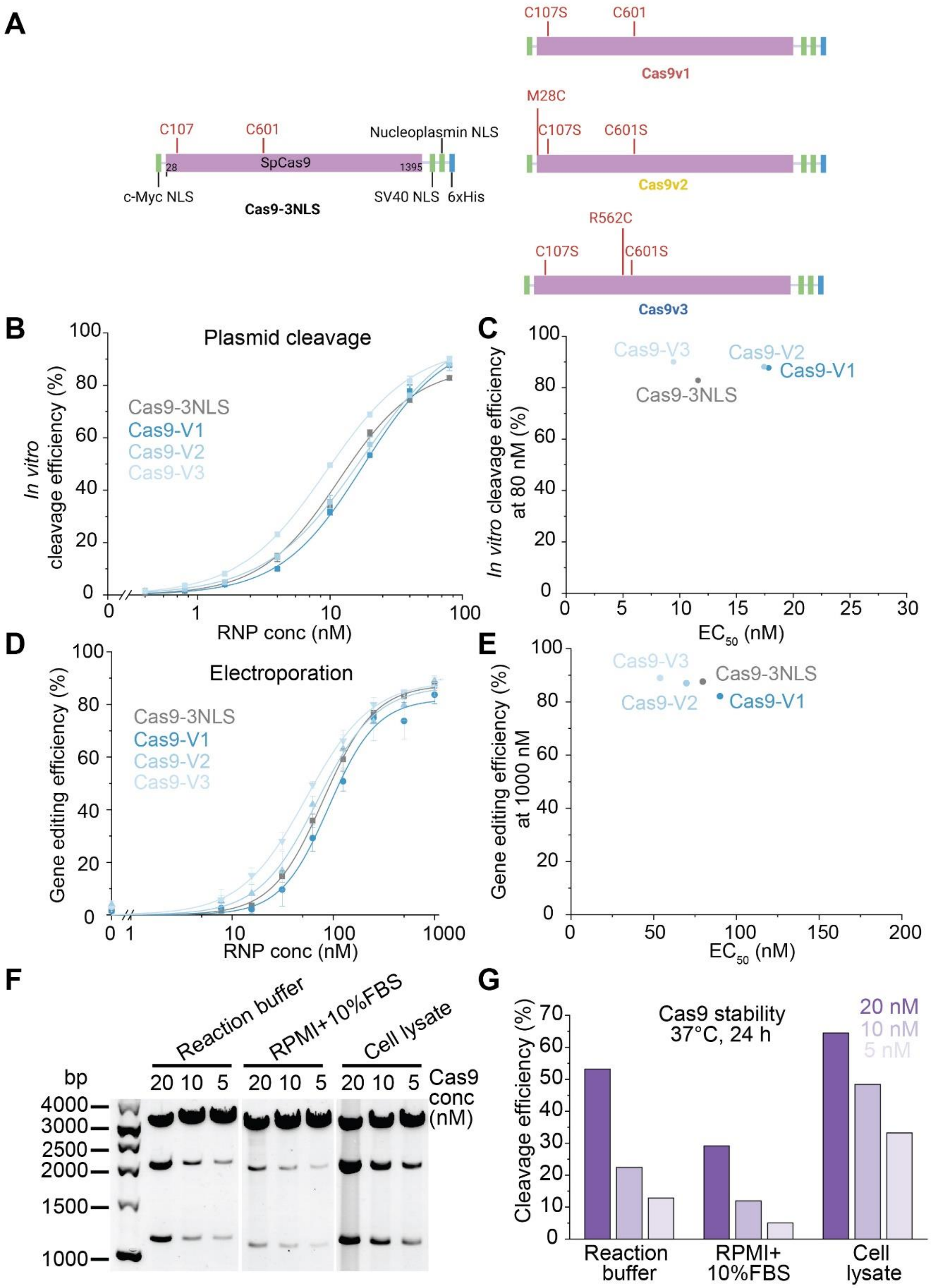
A
C107
C601
Nucleoplasmin NLS
28
SpCas9
1395
c-Myc NLS
SV40 NLS
6xHis
Cas9-3NLS
C107S
C601
Cas9v1
M28C
C107S
C601S
Cas9v2
R562C
C107S
C601S
Cas9v3
B
Plasmid cleavage
In vitro cleavage efficiency (%)
Cas9-3NLS
Cas9-V1
Cas9-V2
Cas9-V3
RNP conc (nM)
C
In vitro cleavage efficiency at 80 nM (%)
EC50 (nM)
D
Electroporation
Gene editing efficiency (%)
RNP conc (nM)
E
Gene editing efficiency (%) at 1000 nM
EC50 (nM)
F
Reaction buffer
RPMI+10%FBS
Cell lysate
bp
20 10 5
Cas9 conc (nM)
4000
3000
2500
2000
1500
1000
G
Cas9 stability 37°C, 24 h
20 nM
10 nM
5 nM
Cleavage efficiency (%)
Reaction buffer
RPMI+ 10%FBS
Cell lysate

**Figure S11. The engineered 3xNLS-SpCas9 variants preserve bioactivity and stability.** (**A**). The design of the three engineered 3xNLS-SpCas9 variants containing single cysteine residue at different Cas9 positions. (**B**) The cleavage efficiency of Cas9-V1, V2 and V3 in comparison with the parent template 3xNLS-SpCas9, assessed in the *in vitro* plasmid cleavage assay. (**C**) The analysis of the $EC_{50}$ and the corresponding cleavage efficiency at 80 nM demonstrated the three engineered 3xNLS-SpCas9 variants retained similar nuclease cleavage activity to the parent 3xNLS-SpCas9 protein. (**D**) The gene editing efficiency of Cas9-V1, V2 and V3 in comparison with the parent template 3xNLS-SpCas9, assessed by elevated mCherry+ levels in flow cytometry 72 hr after the electroporation assay. (**E**) The analysis of the $EC_{50}$ and the corresponding cleavage activity at 1000 nM demonstrated the three engineered 3xNLS-SpCas9 variants retained similar gene editing activity to the parent 3xNLS-SpCas9 protein. (**F**) The stability of the engineered Cas9 (V1 as example) in serum-containing medium and cell lysates. Cas9 was incubated under the indicated conditions at 37 °C for 24 h. The DNA cleavage efficiency was assessed using the *in vitro* plasmid cleavage assay. (**G**) The quantification of the DNA cleavage efficiency indicated the engineered Cas9 retained nuclease activity under the tested solution conditions.

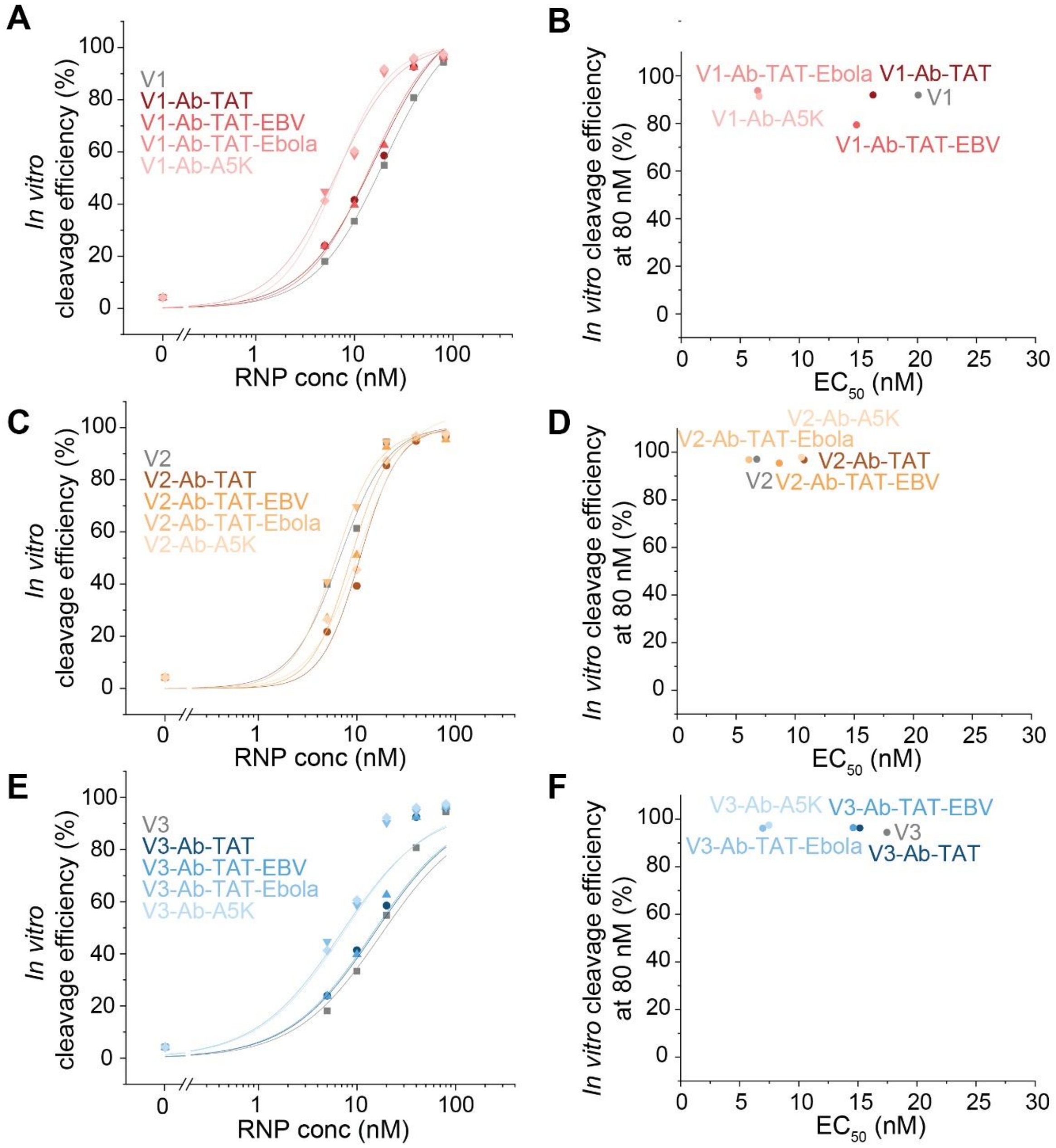


**Figure S12. The assembly of Cas9 tripartite BMCs preserved the precise nuclease activity of Cas9.** (**A, C and E**) The cleavage efficiency of Cas9-Ab tripartite BMCs with TAT, TAT-EBV, TAT-Ebola, A5K as CPPs assessed using the *in vitro* plasmid cleavage assay. All three cas9 variants, V1 (A), V2 (C) and V3 (E), were screened. (**B, D and F**) The analysis of the $EC_{50}$ and the corresponding cleavage activity at 80 nM demonstrated all tested constructs retained the precise nuclease cleavage activity.

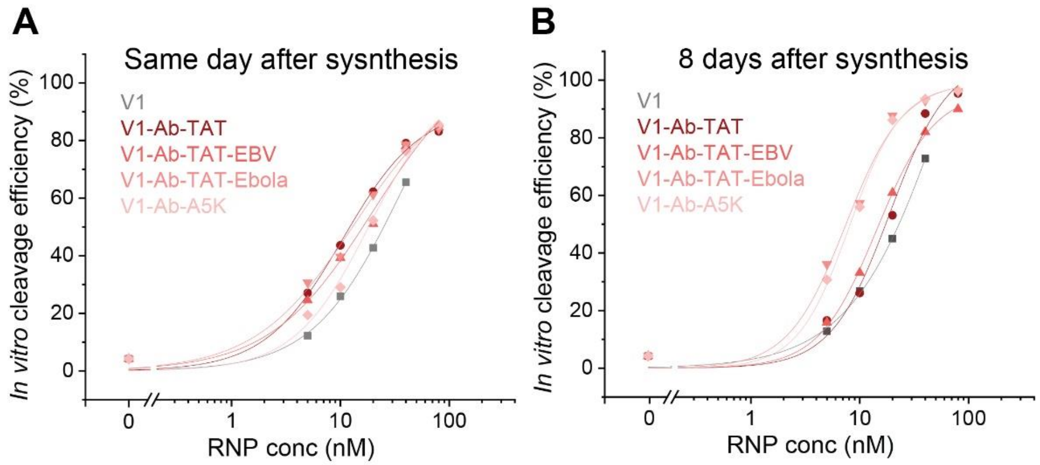


**Figure S13. The stability of the Cas9 tripartite BMCs under storage conditions.** (**A**) The cleavage activity of Cas9 tripartite BMCs assessed the same day after synthesis. (**B**) The cleavage activity of Cas9 tripartite BMCs assessed 8 days after synthesis, samples were stored at 4°C in reaction buffer (50 mM phosphate, 300 mM NaCl, and 50 mM L-arginine hydrochloride, pH 7.0).

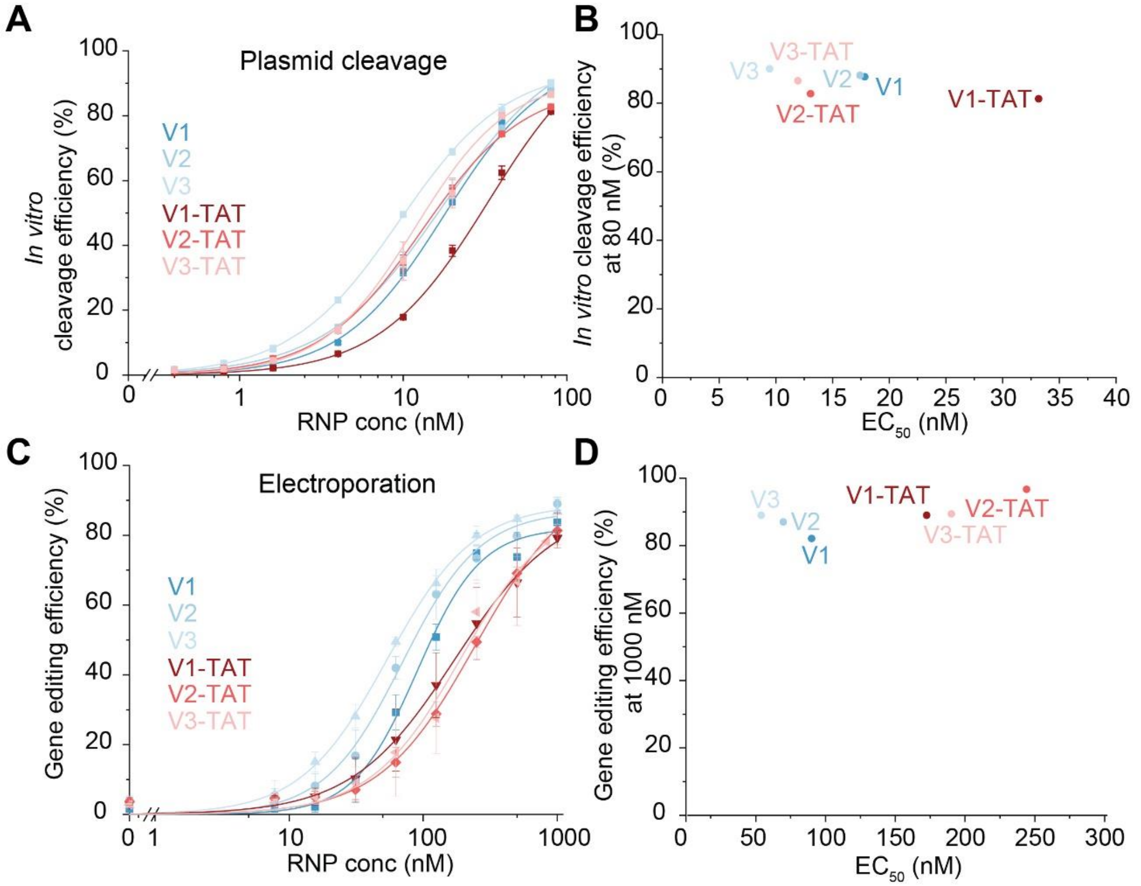


**Figure S14. The modification of Cas9 with positively charged CPP largely preserves its editing activity *in vitro*.** (**A**) The cleavage activity of Cas9-TAT conjugates in comparison with the unconjugated Cas9. All three variants V1, V2 and V3 and the corresponding Cas9-TAT conjugates were assessed in the *in vitro* plasmid cleavage assay. (**B**) The analysis of the $EC_{50}$ and the corresponding cleavage efficiency at 80 nM demonstrated the Cas9-TAT conjugates retained the precise nuclease cleavage function. (**C**) The gene editing activity of Cas9-TAT conjugates in comparison with the unconjugated Cas9. All three variants V1, V2 and V3 and the corresponding Cas9-TAT conjugates were assessed in the electroporation assay. (**D**) The analysis of the $EC_{50}$ and the corresponding cleavage efficiency at 1000 nM demonstrated the Cas9-TAT conjugates retained gene editing activity.

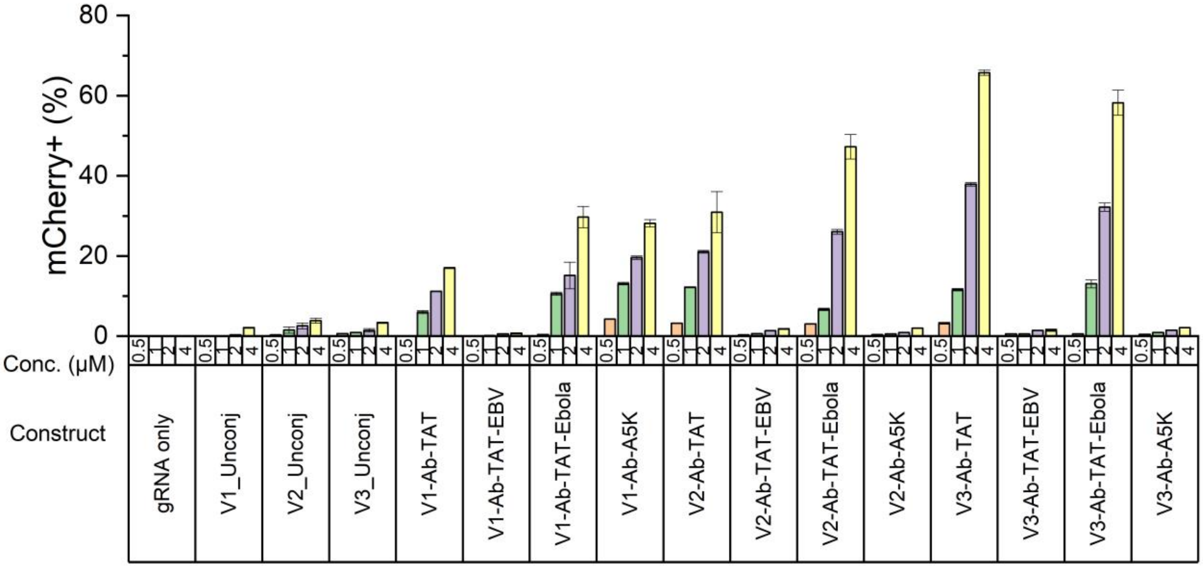


**Figure S15. Cas9 tripartite BMCs functionalized with various CPPs exhibited differential cellular delivery.** While TAT- and TAT-Ebola-modified Cas9 tripartite BMCs showed enhanced delivery and gene editing, Cas9-Ab-TAT-EBV showed minimal editing, and Cas9-Ab-A5K showed enhanced editing only in the V1 variant BMC.

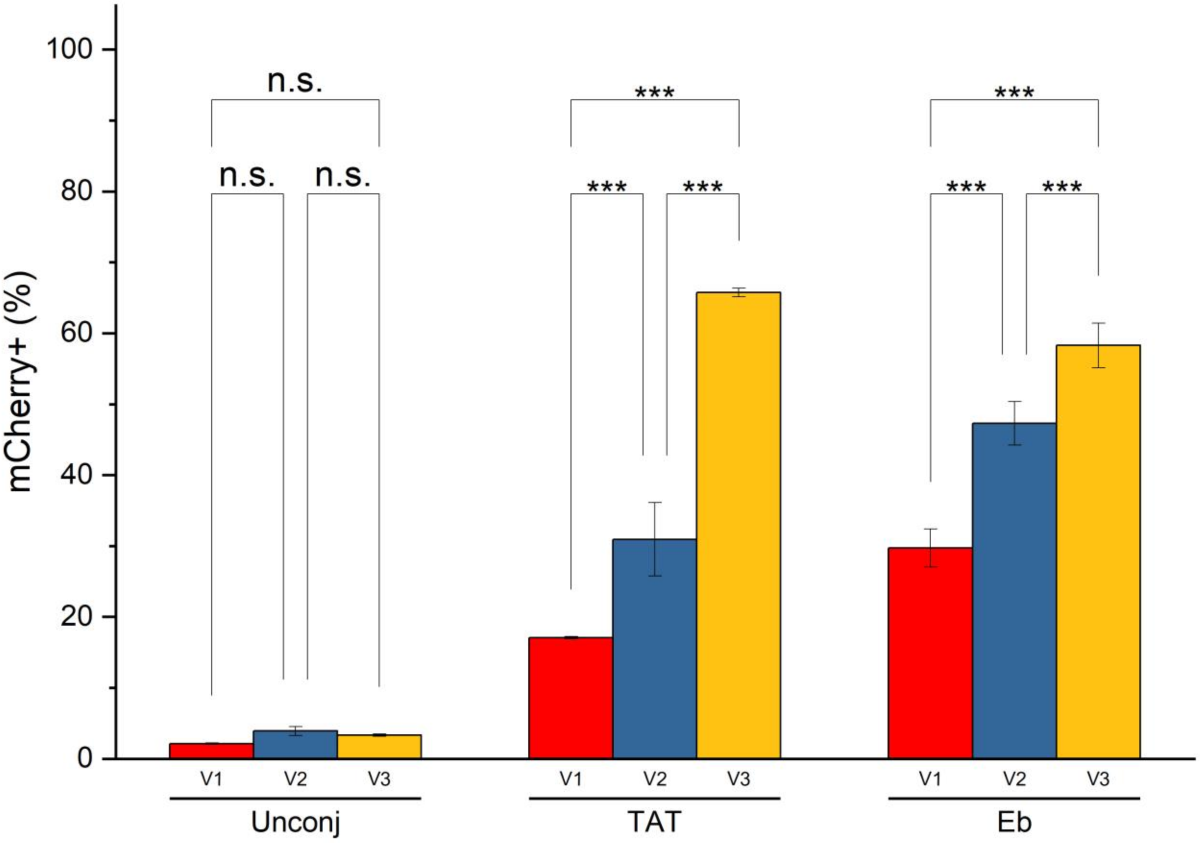


**Figure S16. Effects of different Cas9 variants and CPP conjugation on gene editing activity.** Cas9 tripartite BMC concentration: 4 µM. Statistical significance was estimated using the two-way ANOVA Tukey's test (* $p<=0.05$; ** $p<=0.01$; *** $p<=0.001$).

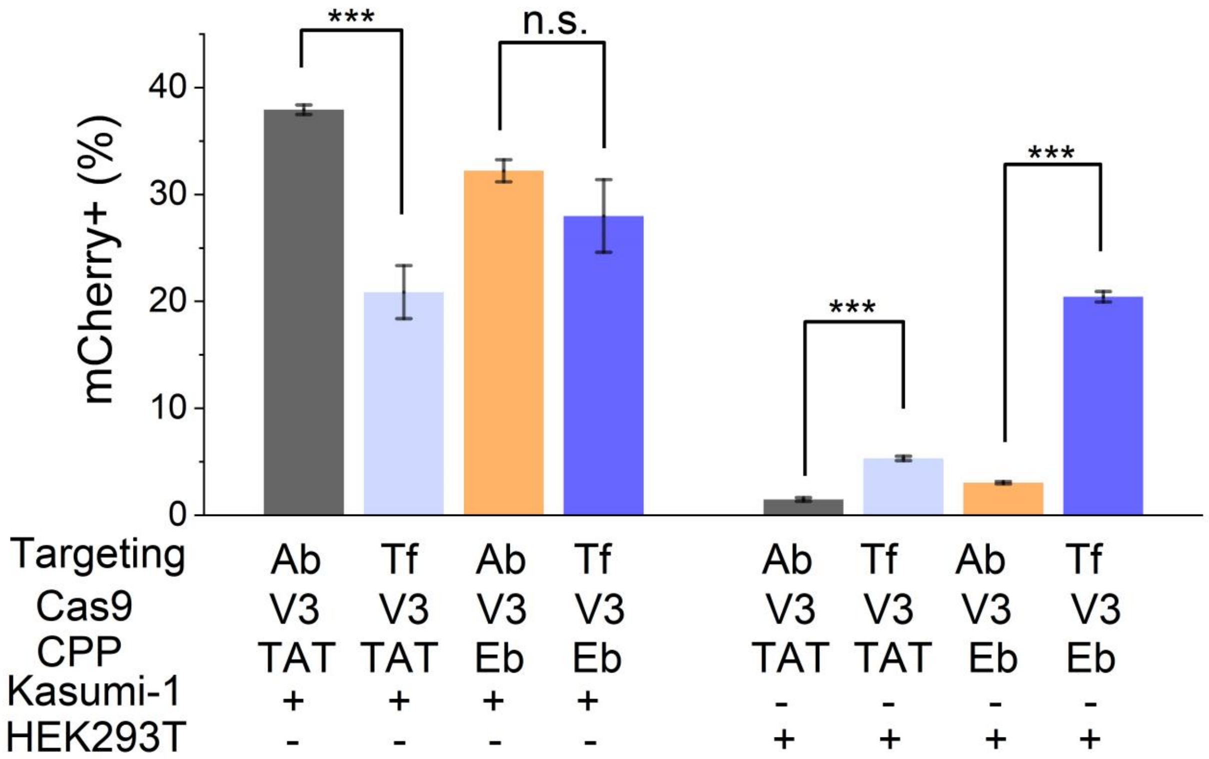


**Figure S17. Cas9 tripartite BMCs were delivered in a cell-specific manner based on the incorporated targeting ligands**. Cas9 tripartite BMC concentration: 2 μM. Statistical significance was estimated using the one-way ANOVA Tukey's test (* $p<=0.05$; ** $p<=0.01$; *** $p<=0.001$).

**Table S1. Molecular details of synthetic peptides used in constructing BMCs**

| Peptide* | Sequence*** | $UV_{280}$ extinction coefficient |
|---|---|---|
| | **Peptidic cargo** | |
| Cys(Npsy)-TAT-FAM | C(Npys)YGRKKRRQRRRK(FAM) | NA |
| CRYBMIM-Cys(Npys) | klenetsmlllelkry[pS]prrslierrGGrrrqrrkkrGyC(Npys)G | NA |
| CRYBMIM-Cys** | klenetsmlllelkry[pS]prrslierrGGrrrqrrkkrGyCG | 2980 |
| | **Tri-functional CPP** | |
| TAT | (DTB)K(azido)YGRKKRRQRRRK(Mal) | 3460 |
| TAT-Ebola | (DTB)K(azido)GAAIGLAWIPYFGPAAYPRKKRRQRRRK(Mal) | 8250 |
| A5K | (DTB)K(azido)GLFEKIEGFIENGWEGMIDGWYGYGRKKRRQRRK(Mal) | 13940 |
| | **Mono-functional CPP** | |
| TAT-EBV | K(azido)IYNGWYAYGRKKRRQRRR | 9530 |
| TAT-Ebola | K(azido)GAAIGLAWIPYFGPAAYPRKKRRQRRR | 8250 |
| A5K | K(azido)GLFEKIEGFIENGWEGMIDGWYGYGRKKRRQRR | 13940 |

*Cys(Npsy)-TAT-FAM was obtained from AnaSpec. All other peptides were synthesized by LifeTein LLC using Fmoc-based solid phase peptide synthesis.
** CRYBMIM-Cys was obtained via reduction of CRYBMIM-Cys(Npys) with TCEP
***Uppercase letter represents L-amino acid, lowercase letter represents D-amino acid.
***Peptides were synthesized with C-terminus amide capping. CRYBMIM-Cys(Npys) was additionally with N-terminus acetyl capping.
***Structures of the modified amino acid were listed below

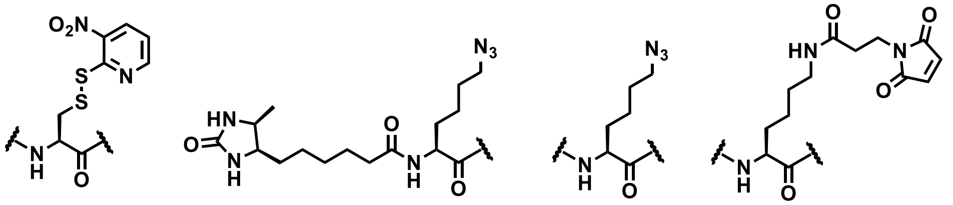

C(Npys)    (DTB)K(azido)    K(azido)    K(Mal)

**Table S2. Molecular details of proteins used in constructing BMCs**

| Protein | Species | MW (Da) | $UV_{280}$ extinction coefficient |
|---|---|---|---|
| 3xNLS-SpCas9 | Streptococcus pyogenes | 166140 | 120575 |
| Cas9-V1 | Streptococcus pyogenes | 166124 | 120450 |
| Cas9-V2 | Streptococcus pyogenes | 166080 | 120450 |
| Cas9-V3 | Streptococcus pyogenes | 166055 | 120450 |
| Transferrin | Homo sapiens | 80000 | 85240 |
| Anti-human c-KIT antibody | Mus musculus | 145000 | 210000 |
| Albumin | Bos taurus | 66400 | 43824 |

### Table S3. Amino acid sequences of the engineered Cas9 variants

cmyc_like_NLS-4aalinker-SpCas9-15aalinker-SV40_NLS-4aalinker-nucleoplasmin_NLS-6XHis_tag

| Variant name | Cys position | Engineered site |
|---|---|---|
| 3xNLS-SpCas9 | C107, C601 | NA |

MASMTGGQQMGRAAPAAKKKKLDGSVDMDKKYSIGLDIGTNSVGWAVITDEYKVPSKKFKVLGNTDRHSIKKNLIGALLF
DSGETAEATRLKRTARRRYTRRKNRICYLQEIFSNEMAKVDDSFFHRLEESFLVEEDKKHERHPIFGNIVDEVAYHEKYPTIYH
LRKKLVDSTDKADLRLIYLALAHMIKFRGHFLIEGDLNPDNSDVDKLFIQLVQTYNQLFEENPINASGVDAKAILSARLSKSR
RLENLIAQLPGEKKNGLFGNLIALSLGLTPNFKSNFDLAEDAKLQLSKDTYDDDLDNLLAQIGDQYADLFLAAKNLSDAILLS
DILRVNTEITKAPLSASMIKRYDEHHQDLTLLKALVRQQLPEKYKEIFFDQSKNGYAGYIDGGASQEEFYKFIKPILEKMDGTE
ELLVKLNREDLLRKQRTFDNGSIPHQIHLGELHAILRRQEDFYPFLKDNREKIEKILTFRIPYYVGPLARGNSRFAWMTRKSEE
TITPWNFEEVVDKGASAQSFIERMTNFDKNLPNEKVLPKHSLLYEYFTVYNELTKVKYVTEGMRKPAFLSGEQKKAIVDLLF
KTNRKVTVKQLKEDYFKKIECFDSVEISGVEDRFNASLGTYHDLLKIIKDKDFLDNEENEDILEDIVLTLTLFEDREMIEERLK
TYAHLFDDKVMKQLKRRRYTGWGRLSRKLINGIRDKQSGKTILDFLKSDGFANRNFMQLIHDDSLTFKEDIQKAQVSGQGD
SLHEHIANLAGSPAIKKGILQTVKVVDELVKVMGRHKPENIVIEMARENQTTQKGQKNSRERMKRIEEGIKELGSQILKEHPV
ENTQLQNEKLYLYYLQNGRDMYVDQELDINRLSDYDVDHIVPQSFLKDDSIDNKVLTRSDKNRGKSDNVPSEEVVKKMKN
YWRQLLNAKLITQRKFDNLTKAERGGLSELDKAGFIKRQLVETRQITKHVAQILDSRMNTKYDENDKLIREVKVITLKSKLVS
DFRKDFQFYKVREINNYHHAHDAYLNAVVGTALIKKYPKLESEFVYGDYKVYDVRKMIAKSEQEIGKATAKYFFYSNIMNF
FKTEITLANGEIRKRPLIETNGETGEIVWDKGRDFATVRKVLSMPQVNIVKKTEVQTGGFSKESILPKRNSDKLIARKKDWDP
KKYGGFDSPTVAYSVLVVAKVEKGKSKKLKSVKELLGITIMERSSFEKNPIDFLEAKGYKEVKKDLIIKLPKYSLFELENGRK
RMLASAGELQKGNELALPSKYVNFLYLASHYEKLKGSPEDNEQKQLFVEQHKHYLDEIIEQISEFSKRVILADANLDKVLSA
YNKHRDKPIREQAENIIHLFTLTNLGAPAAFKYFDTTIDRKRYTSTKEVLDATLIHQSITGLYETRIDLSQLGGD
TGGGPGGGAAAGSGSPKKKRKVGSGSKRPAATKKAGQAKKKKLEHHHHHH

| Variant name | Cys position | Engineered site |
|---|---|---|
| Cas9-V1 | C601 | C107S |

MASMTGGQQMGRAAPAAKKKKLDGSVDMDKKYSIGLDIGTNSVGWAVITDEYKVPSKKFKVLGNTDRHSIKKNLIGALLF
DSGETAEATRLKRTARRRYTRRKNRISYLQEIFSNEMAKVDDSFFHRLEESFLVEEDKKHERHPIFGNIVDEVAYHEKYPTIYH
LRKKLVDSTDKADLRLIYLALAHMIKFRGHFLIEGDLNPDNSDVDKLFIQLVQTYNQLFEENPINASGVDAKAILSARLSKSR
RLENLIAQLPGEKKNGLFGNLIALSLGLTPNFKSNFDLAEDAKLQLSKDTYDDDLDNLLAQIGDQYADLFLAAKNLSDAILLS
DILRVNTEITKAPLSASMIKRYDEHHQDLTLLKALVRQQLPEKYKEIFFDQSKNGYAGYIDGGASQEEFYKFIKPILEKMDGTE
ELLVKLNREDLLRKQRTFDNGSIPHQIHLGELHAILRRQEDFYPFLKDNREKIEKILTFRIPYYVGPLARGNSRFAWMTRKSEE
TITPWNFEEVVDKGASAQSFIERMTNFDKNLPNEKVLPKHSLLYEYFTVYNELTKVKYVTEGMRKPAFLSGEQKKAIVDLLF
KTNRKVTVKQLKEDYFKKIECFDSVEISGVEDRFNASLGTYHDLLKIIKDKDFLDNEENEDILEDIVLTLTLFEDREMIEERLK
TYAHLFDDKVMKQLKRRRYTGWGRLSRKLINGIRDKQSGKTILDFLKSDGFANRNFMQLIHDDSLTFKEDIQKAQVSGQGD
SLHEHIANLAGSPAIKKGILQTVKVVDELVKVMGRHKPENIVIEMARENQTTQKGQKNSRERMKRIEEGIKELGSQILKEHPV
ENTQLQNEKLYLYYLQNGRDMYVDQELDINRLSDYDVDHIVPQSFLKDDSIDNKVLTRSDKNRGKSDNVPSEEVVKKMKN
YWRQLLNAKLITQRKFDNLTKAERGGLSELDKAGFIKRQLVETRQITKHVAQILDSRMNTKYDENDKLIREVKVITLKSKLVS
DFRKDFQFYKVREINNYHHAHDAYLNAVVGTALIKKYPKLESEFVYGDYKVYDVRKMIAKSEQEIGKATAKYFFYSNIMNF
FKTEITLANGEIRKRPLIETNGETGEIVWDKGRDFATVRKVLSMPQVNIVKKTEVQTGGFSKESILPKRNSDKLIARKKDWDP
KKYGGFDSPTVAYSVLVVAKVEKGKSKKLKSVKELLGITIMERSSFEKNPIDFLEAKGYKEVKKDLIIKLPKYSLFELENGRK
RMLASAGELQKGNELALPSKYVNFLYLASHYEKLKGSPEDNEQKQLFVEQHKHYLDEIIEQISEFSKRVILADANLDKVLSA
YNKHRDKPIREQAENIIHLFTLTNLGAPAAFKYFDTTIDRKRYTSTKEVLDATLIHQSITGLYETRIDLSQLGGDTGGGPGGGA
AAGSGSPKKKRKVGSGSKRPAATKKAGQAKKKKLEHHHHHH

| Variant name | Cys position | Engineered site |
|---|---|---|
| Cas9-V2 | M28C | C107S, C601S |

MASMTGGQQMGRAAPAAKKKKLDGSVDCDKKYSIGLDIGTNSVGWAVITDEYKVPSKKFKVLGNTDRHSIKKNLIGALLF
DSGETAEATRLKRTARRRYTRRKNRISYLQEIFSNEMAKVDDSFFHRLEESFLVEEDKKHERHPIFGNIVDEVAYHEKYPTIYH
LRKKLVDSTDKADLRLIYLALAHMIKFRGHFLIEGDLNPDNSDVDKLFIQLVQTYNQLFEENPINASGVDAKAILSARLSKSR
RLENLIAQLPGEKKNGLFGNLIALSLGLTPNFKSNFDLAEDAKLQLSKDTYDDDLDNLLAQIGDQYADLFLAAKNLSDAILLS
DILRVNTEITKAPLSASMIKRYDEHHQDLTLLKALVRQQLPEKYKEIFFDQSKNGYAGYIDGGASQEEFYKFIKPILEKMDGTE
ELLVKLNREDLLRKQRTFDNGSIPHQIHLGELHAILRRQEDFYPFLKDNREKIEKILTFRIPYYVGPLARGNSRFAWMTRKSEE
TITPWNFEEVVDKGASAQSFIERMTNFDKNLPNEKVLPKHSLLYEYFTVYNELTKVKYVTEGMRKPAFLSGEQKKAIVDLLF
KTNRKVTVKQLKEDYFKKIESFDSVEISGVEDRFNASLGTYHDLLKIIKDKDFLDNEENEDILEDIVLTLTLFEDREMIEERLK
TYAHLFDDKVMKQLKRRRYTGWGRLSRKLINGIRDKQSGKTILDFLKSDGFANRNFMQLIHDDSLTFKEDIQKAQVSGQGD
SLHEHIANLAGSPAIKKGILQTVKVVDELVKVMGRHKPENIVIEMARENQTTQKGQKNSRERMKRIEEGIKELGSQILKEHPV
ENTQLQNEKLYLYYLQNGRDMYVDQELDINRLSDYDVDHIVPQSFLKDDSIDNKVLTRSDKNRGKSDNVPSEEVVKKMKN
YWRQLLNAKLITQRKFDNLTKAERGGLSELDKAGFIKRQLVETRQITKHVAQILDSRMNTKYDENDKLIREVKVITLKSKLVS
DFRKDFQFYKVREINNYHHAHDAYLNAVVGTALIKKYPKLESEFVYGDYKVYDVRKMIAKSEQEIGKATAKYFFYSNIMNF
FKTEITLANGEIRKRPLIETNGETGEIVWDKGRDFATVRKVLSMPQVNIVKKTEVQTGGFSKESILPKRNSDKLIARKKDWDP
KKYGGFDSPTVAYSVLVVAKVEKGKSKKLKSVKELLGITIMERSSFEKNPIDFLEAKGYKEVKKDLIIKLPKYSLFELENGRK
RMLASAGELQKGNELALPSKYVNFLYLASHYEKLKGSPEDNEQKQLFVEQHKHYLDEIIEQISEFSKRVILADANLDKVLSA
YNKHRDKPIREQAENIIHLFTLTNLGAPAAFKYFDTTIDRKRYTSTKEVLDATLIHQSITGLYETRIDLSQLGGDTGGGPGGGA
AAGSGSPKKKRKVGSGSKRPAATKKAGQAKKKKLEHHHHHH

| Variant name | Cys position | Engineered site |
|---|---|---|
| Cas9-V3 | R562C | C107S, C601S |

MASMTGGQQMGRAAPAAKKKKLDGSVDMDKKYSIGLDIGTNSVGWAVITDEYKVPSKKFKVLGNTDRHSIKKNLIGALLF
DSGETAEATRLKRTARRRYTRRKNRISYLQEIFSNEMAKVDDSFFHRLEESFLVEEDKKHERHPIFGNIVDEVAYHEKYPTIYH
LRKKLVDSTDKADLRLIYLALAHMIKFRGHFLIEGDLNPDNSDVDKLFIQLVQTYNQLFEENPINASGVDAKAILSARLSKSR
RLENLIAQLPGEKKNGLFGNLIALSLGLTPNFKSNFDLAEDAKLQLSKDTYDDDLDNLLAQIGDQYADLFLAAKNLSDAILLS
DILRVNTEITKAPLSASMIKRYDEHHQDLTLLKALVRQQLPEKYKEIFFDQSKNGYAGYIDGGASQEEFYKFIKPILEKMDGTE
ELLVKLNREDLLRKQRTFDNGSIPHQIHLGELHAILRRQEDFYPFLKDNREKIEKILTFRIPYYVGPLARGNSRFAWMTRKSEE
TITPWNFEEVVDKGASAQSFIERMTNFDKNLPNEKVLPKHSLLYEYFTVYNELTKVKYVTEGMCKPAFLSGEQKKAIVDLLF
KTNRKVTVKQLKEDYFKKIESFDSVEISGVEDRFNASLGTYHDLLKIIKDKDFLDNEENEDILEDIVLTLTLFEDREMIEERLK
TYAHLFDDKVMKQLKRRRYTGWGRLSRKLINGIRDKQSGKTILDFLKSDGFANRNFMQLIHDDSLTFKEDIQKAQVSGQGD
SLHEHIANLAGSPAIKKGILQTVKVVDELVKVMGRHKPENIVIEMARENQTTQKGQKNSRERMKRIEEGIKELGSQILKEHPV
ENTQLQNEKLYLYYLQNGRDMYVDQELDINRLSDYDVDHIVPQSFLKDDSIDNKVLTRSDKNRGKSDNVPSEEVVKKMKN
YWRQLLNAKLITQRKFDNLTKAERGGLSELDKAGFIKRQLVETRQITKHVAQILDSRMNTKYDENDKLIREVKVITLKSKLVS
DFRKDFQFYKVREINNYHHAHDAYLNAVVGTALIKKYPKLESEFVYGDYKVYDVRKMIAKSEQEIGKATAKYFFYSNIMNF
FKTEITLANGEIRKRPLIETNGETGEIVWDKGRDFATVRKVLSMPQVNIVKKTEVQTGGFSKESILPKRNSDKLIARKKDWDP
KKYGGFDSPTVAYSVLVVAKVEKGKSKKLKSVKELLGITIMERSSFEKNPIDFLEAKGYKEVKKDLIIKLPKYSLFELENGRK
RMLASAGELQKGNELALPSKYVNFLYLASHYEKLKGSPEDNEQKQLFVEQHKHYLDEIIEQISEFSKRVILADANLDKVLSA
YNKHRDKPIREQAENIIHLFTLTNLGAPAAFKYFDTTIDRKRYTSTKEVLDATLIHQSITGLYETRIDLSQLGGDTGGGPGGGA
AAGSGSPKKKRKVGSGSKRPAATKKAGQAKKKKLEHHHHHH

**Table S4.** $EC_{50}$ values for the Cas9 tripartite BMCs in the plasmid cleavage assay

| Construct | $EC_{50}$ (nM) | *Fold change to Unconjugated Cas9 |
|---|---|---|
| V1_Unconj | 18.1 | 1.0 |
| V2_Unconj | 15.2 | 1.0 |
| V3_Unconj | 9.5 | 1.0 |
| V1-Tf-TAT | 24.6 | 1.4 |
| V2-Tf-TAT | 25.1 | 1.7 |
| V3-Tf-TAT | 19.1 | 2.0 |
| V1-Ab-TAT | 24.1 | 1.3 |
| V2-Ab-TAT | 20.3 | 1.3 |
| V3-Ab-TAT | 16.7 | 1.8 |

*The fold changes are calculated based on the $EC_{50}$ values of corresponding Cas9 variants (V1/V2/V3).

**Table S5.** $EC_{50}$ values for the Cas9 tripartite BMCs in the plasmid cleavage assay

| Construct | $EC_{50}$ (nM) | *Fold change to Unconjugated Cas9 |
|---|---|---|
| Unconj_Cas9-V1 | 20 | 1.0 |
| Cas9-V1-Ab-TAT | 16 | 0.8 |
| Cas9-V1-Ab-TAT-EBV | 15 | 0.7 |
| Cas9-V1-Ab-TAT-Ebola | 6.5 | 0.3 |
| Cas9-V1-Ab-A5K | 6.6 | 0.3 |
| Unconj_Cas9-V2 | 6.8 | 1.0 |
| Cas9-V2-Ab-TAT | 11 | 1.6 |
| Cas9-V2-Ab-TAT-EBV | 8.7 | 1.3 |
| Cas9-V2-Ab-TAT-Ebola | 6.1 | 0.9 |
| Cas9-V2-Ab-A5K | 11 | 1.6 |
| Unconj_Cas9-V3 | 18 | 1.0 |
| Cas9-V3-Ab-TAT | 15 | 0.9 |
| Cas9-V3-Ab-TAT-EBV | 15 | 0.8 |
| Cas9-V3-Ab-TAT-Ebola | 7.0 | 0.4 |
| Cas9-V3-Ab-A5K | 7.5 | 0.4 |

*The fold changes are calculated based on the $EC_{50}$ values of corresponding Cas9 variants (V1/V2/V3).

**Table S6.** $EC_{50}$ values for the Cas9 tripartite BMCs in the cellular electroporation experiment

| Construct | $EC_{50}$ (nM) | *Fold change to Unconjugated Cas9 |
|---|---|---|
| Unconj_Cas9-V1 | 60.0 | 1.0 |
| Cas9-V1-Ab-TAT | 76.1 | 1.3 |
| Cas9-V1-Ab-TAT-EBV | 81.3 | 1.4 |
| Cas9-V1-Ab-TAT-Ebola | 62.5 | 1.0 |
| Cas9-V1-Ab-A5K | 48.0 | 0.8 |
| Unconj_Cas9-V2 | 34.0 | 1.0 |
| Cas9-V2-Ab-TAT | 48.6 | 1.4 |
| Cas9-V2-Ab-TAT-EBV | 43.2 | 1.3 |
| Cas9-V2-Ab-TAT-Ebola | 47.3 | 1.4 |
| Cas9-V2-Ab-A5K | 28.4 | 0.8 |
| Unconj_Cas9-V3 | 48.1 | 1.0 |
| Cas9-V3-Ab-TAT | 127 | 2.6 |
| Cas9-V3-Ab-TAT-EBV | 85.1 | 1.8 |
| Cas9-V3-Ab-TAT-Ebola | 110 | 2.3 |
| Cas9-V3-Ab-A5K | 73.7 | 1.5 |

*The fold changes are calculated based on the $EC_{50}$ values of corresponding Cas9 variants (V1/V2/V3).
